# Selenocysteine metabolism is a targetable vulnerability in *MYCN*-amplified cancers

**DOI:** 10.1101/2022.05.17.492172

**Authors:** Hamed Alborzinia, Zhiyi Chen, Umut Yildiz, Florencio Porto Freitas, Felix C.E. Vogel, Julianna Varga, Jasmin Batani, Christoph Bartenhagen, Werner Schmitz, Gabriele Büchel, Bernhard Michalke, Jashuo Zheng, Svenja Meierjohann, Enrico Girardi, Elisa Espinet, Andres Florez, Ancely Ferreira dos Santos, Nesrine Aroua, Lisa Schlicker, Thamara N. Xavier da Silva, Adriana Przybylla, Petra Zeisberger, Giulio Superti-Furga, Martin Eilers, Marcus Conrad, Matthias Fischer, Almut Schulze, Andreas Trumpp, José Pedro Friedmann Angeli

## Abstract

Understanding the operational molecular, and metabolic networks that determine the balance between pro- and anti-ferroptotic regulatory pathways could unravel unique vulnerabilities to be exploited for cancer therapy. Here we identify the selenoprotein P (SELENOP) receptor, LRP8, as a key determinant protecting MYCN-amplified neuroblastoma cells from ferroptosis in vitro and in orthotopic neuroblastoma mouse models. Specifically, the exquisite dependency on LRP8-mediated selenocysteine import is caused by the failure of MYCN-amplified cells to efficiently utilize alternative forms of selenium/selenocysteine based uptake necessary for selenoprotein biosynthesis. Increased activity of one of such transporters, SLC7A11, in MYCN-amplified cells leads to cysteine overload, progressive mitochondrial decline and impaired proliferation. These data reveal in LRP8 a targetable, and specific vulnerability of MYCN-amplified neuroblastoma cells and disclose a yet-unaccounted mechanism for selective ferroptosis induction that has the potential to become an important therapeutic entry point for MYCN-amplified neuroblastoma.

**Statement of significance:** Given the largely unsuccessful repurposing of adult oncology drugs for the treatment of neuroblastoma, our discoveries pave the way for novel ferroptosis based strategies for this entity. Specifically, targeting of LRP8 may offer novel therapeutic and safer opportunities for a number of pediatric malignancies and MYCN driven cancers.

## Introduction

Ferroptosis is a unique cell death modality that is attracting increasing interest as a means to eradicate therapeutically challenging tumor entities (*1, 2*). At the molecular level, ferroptosis has been shown to be primarily suppressed by the selenoprotein glutathione peroxidase 4 (GPX4) (*3*). GPX4 utilizes reducing equivalents from glutathione (GSH) to suppress the accumulation of lipid hydroperoxides, and, ultimately, the induction of cell death (*4*). This central role of GPX4 has spurred an intense search for strategies and molecular tools able to inhibit its activity (*2, 3, 5, 6*). Nonetheless, the lack of suitable *in vivo* GPX4 inhibitors and the foreseen systemic toxicity of such compounds (*7-9*) are still limiting and pose cautionary aspects for the translation of these discoveries into anti-cancer therapies. Recently we, and others, have uncovered that high-risk *MYCN*-amplified neuroblastomas are characterized by a striking GPX4 dependency (*10-12*). The molecular determinants of this increased dependency remain elusive, and untangling the mechanisms dictating ferroptosis hypersensitive states could pave the way to specifically exploit these vulnerabilities. In the present work, using genome-wide and single-cell transcriptomics CRISPR-activation (CRISPRa) screens, we identify novel regulators of ferroptosis and detect shared transcriptional signatures and states regulating ferroptosis hypersensitivity. Specifically, we identify the low-density lipoprotein receptor (LDLR) related protein 8 (LRP8, also known as APOER2) as a critical bottleneck in selenium/selenocysteine metabolism in *MYCN*-amplified entities. We found that the MYCN-associated vulnerability is due to the incompatibility of *MYCN*-amplified cells to activate alternative selenium/selenocysteine pathways, such as SLC7A11, required to support selenoprotein translation. Therefore, our work demonstrates that different pathways of selenium/selenocysteine acquisition are associated with unique metabolic consequences that can be accompanied by a severe metabolic disruption in specific oncogenic contexts. The identification of these metabolic vulnerabilities offers unanticipated opportunities to specifically, selectively and safely target LRP8 to induce ferroptosis for therapeutic benefit.

## Results

### CRISPR activation screens identify novel regulator of ferroptosis

In order to gain additional insight into the process of ferroptosis, we reasoned that understanding the mechanisms capable of suppressing ferroptosis in intrinsically hypersensitive cells would provide a means to identify as yet uncharacterized regulators of ferroptosis. To this end, we initially analyzed data from the depmap portal (www.depmap.org) in search for cell lines that are hypersensitive to the GPX4 inhibitors RSL3 and ML210 (**Fig. 1A**). We selected the *MYCN*-amplified neuroblastoma cell line SK-N-DZ, a representative of tumor entities that still defy current treatments and for which we and others have previously reported a marked dependency on GPX4 (*10-13*), thus providing an ideal setting to interrogate the mechanisms underlying this hypersensitivity. Using this cell line, we performed genome-wide CRISPR activation (CRISPRa) screens during which we induced ferroptosis via two non-overlapping mechanisms (**Fig. 1B)**, namely via i) GPX4 and via ii) system Xc-inhibition. Screen deconvolution allowed us to identify several known and novel ferroptosis regulators (**Fig. 1C**). A secondary screen focusing on the obtained hits confirmed the resistance phenotype for the majority of the scoring hits when selected against the GPX4 inhibitor RSL3 (**Fig. 1D**). In order to provide an unbiased understanding of the differential mechanism of cellular states that prevent ferroptosis, we coupled the focused screen to single-cell RNA-seq (scRNA-seq, CROP-seq) as a readout (**Fig. 1E**). The focused library consisted of two guide RNAs (gRNAs) targeting each of the selected 36 candidate genes together with ten non-targeting control gRNAs (NT ctrl). We obtained high-quality data from ∼11,000 CRISPRa-assigned cells (mean of ∼78,000 reads and median of ∼6,600 genes per cell) (fig. S1A, B). Selecting cells with a strong CRISPRa phenotype (see methods) allowed us to retrieve 130 cells on average for each of the selected hits (32/36 target gene identities detected, fig. S1B). Target identities for which cells could not be assigned (*SLC7A11, TAF4, ARR3, SLC13A4*) likely conferred a negative impact on cell proliferation as a consequence of target gene overexpression in the absence of ferroptosis-inducing agents (fig. S1B). As expected, cells assigned to a particular perturbation cluster showed an increased expression of the targeted gene (**Fig. 1F** and fig. S1D, E). Interestingly, the impact of several CRISPRa phenotypes converged on the expression of known ferroptosis regulators. For instance, overexpression phenotypes of *IRS4* and *MET* shared the upregulation of known ferroptosis suppressors, including the transcription factor *NFE2L2* that drives the expression of genes involved in redox signalling and oxidative protection (*14*), the heat-shock protein *HSPB1*, as well as the CoQ oxidoreductases *AIFM2* (also known as *FSP1* (*15, 16*)) (fig. S1E, F). Gene Ontology analysis of differentially expressed genes upon CRISPRa showed significant enrichment in processes involved in cellular detoxification, energy metabolism, and proteasomal degradation that have all been previously associated with ferroptosis (*17-19*) (fig. S1G). Interestingly, the highest scoring hit in our primary screen, *LRP8*, clustered together with the *GPX4* overexpression phenotype in our scCRISPRa screen, underscoring that the mechanism underlying ferroptosis resistance might overlap and be shared between multiple groups (**Fig. 1F** and fig. S1C, E). Thus, our scCRISPRa secondary screen faithfully recapitulated the gRNA identity in single-cell assays and simultaneously provided transcriptional signatures of different ferroptosis resistant states derived from the genome-wide screen.

**Fig. 1.**
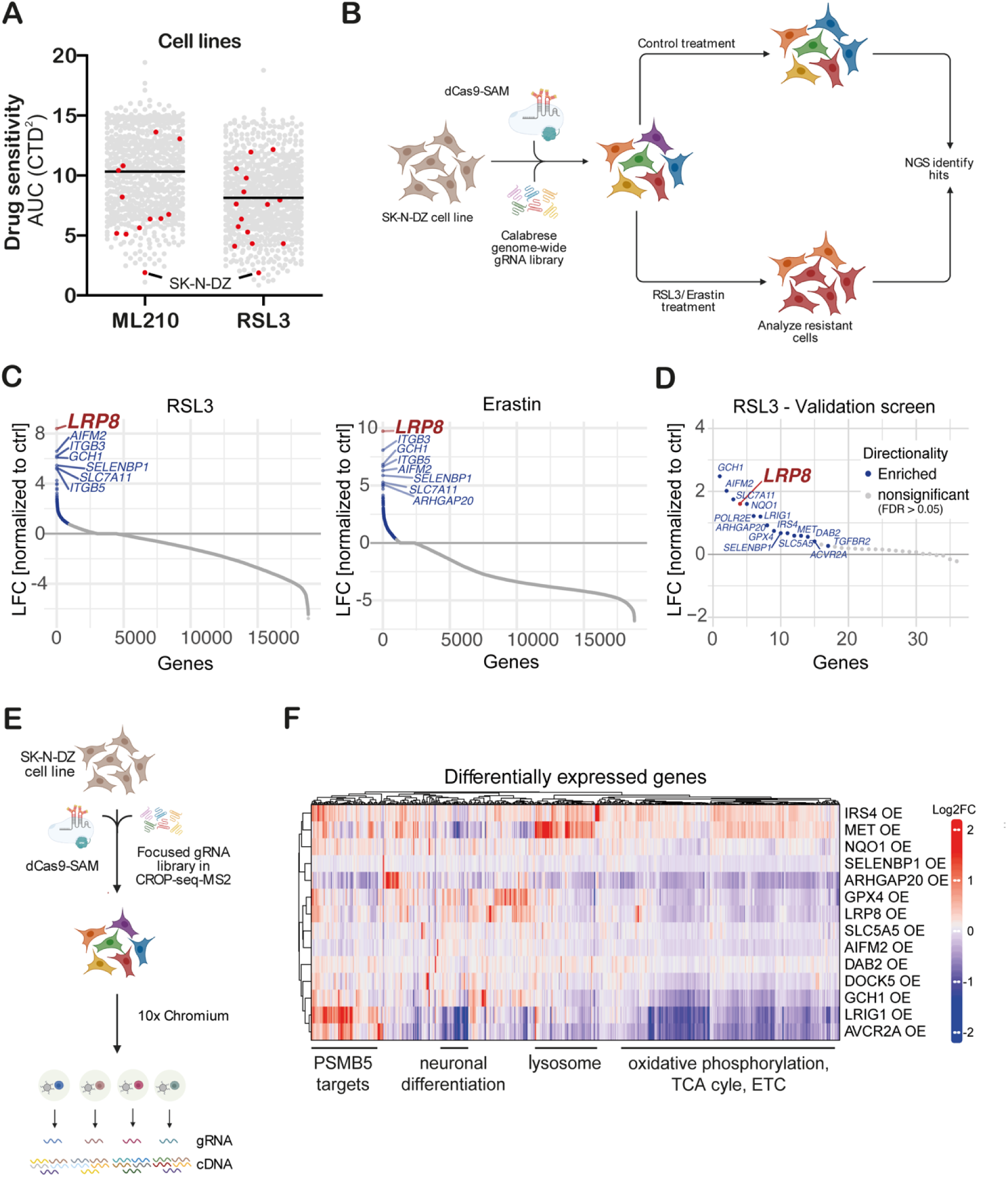
Genome-wide CRISPR activation screen identifies negative regulators of ferroptosis. **(A)** Analysis of the depmap portal (www.depmap.org) reveals *MYCN*-amplified SK-N-DZ as hypersensitive cell lines to the GPX4 inhibitors RSL3 and ML210. (**B)** Strategy of the genome-wide CRISPR activation (CRISPRa) screen in *MYCN-*amplified SK-N-DZ cells. (**C)** Overexpression phenotypes conferring resistance to 300 nM RSL3 (left) or 1 μM Erastin (right) treatment. Significant hits are marked in blue (FDR ≤ 0.05), while the highest scoring hit, *LRP8*, is highlighted in red. (**D)** Overexpression phenotypes conferring resistance to RSL3 (100 nM) induced ferroptosis in the pooled validation CRISPRa screen. Significantly enriched genes are marked and labelled in blue (FDR ≤ 0.05). The highest scoring hit from the primary screens, *LRP8*, is highlighted in red (**E)** Strategy of the single-cell CRISPRa screen to characterize hits from the ferroptosis-resistance screen. Guide RNA labels are recovered alongside the whole transcriptome readout for each cell. **(F)** Transcriptomic consequences of CRISPRa of 14 scoring hits from the primary and the validation screens. Each row represents one CRISPRa cluster. For each cluster, the top 50 genes with the most significantly differential expression (compared to the non-targeting control cluster) were selected and merged to a signature gene list represented by the columns. Columns and rows were hierarchically clustered based on Pearson correlation.

### SELENOP is a critical source of selenium to support the growth of a subset of MYCN-amplified neuroblastomas

The identification of LRP8 as a critical regulator of ferroptosis sensitivity is in agreement with data from the cancer therapeutic response portal (CTRP), showing that expression of *LRP8* strongly correlates with resistance to GPX4 inhibitors (ML210, RSL3 and ML162) (fig. S2A) (*20*). To functionally validate our findings, we first overexpressed *LRP8* using a CRISPRa system in which SK-N-DZ cells became robustly resistant to RSL3 treatment (fig. S2B). Similarly, stable overexpression of a Flag-tagged h*LRP8* in SK-N-DZ cells conveyed resistance to a panel of ferroptosis inducers covering differential modes of action (**Fig. 2A, B**). In line with a specific inhibitory role for LRP8 in ferroptosis, an overall impact on sensitivity towards other cytotoxic compounds was not observed (fig. S2C). In accordance with the protective effect observed, *LRP8* overexpression suppressed lipid peroxidation in cells treated with a GPX4 inhibitor (**Fig. 2C**). Furthermore, overexpression of *LRP8* was able to increase ferroptosis resistance in a larger panel of cell lines (**Fig. 2D**). No other members of the LRP family showed similar effects when overexpressed, highlighting the specific role of LRP8 (fig. S2D, E).

**Fig. 2.**
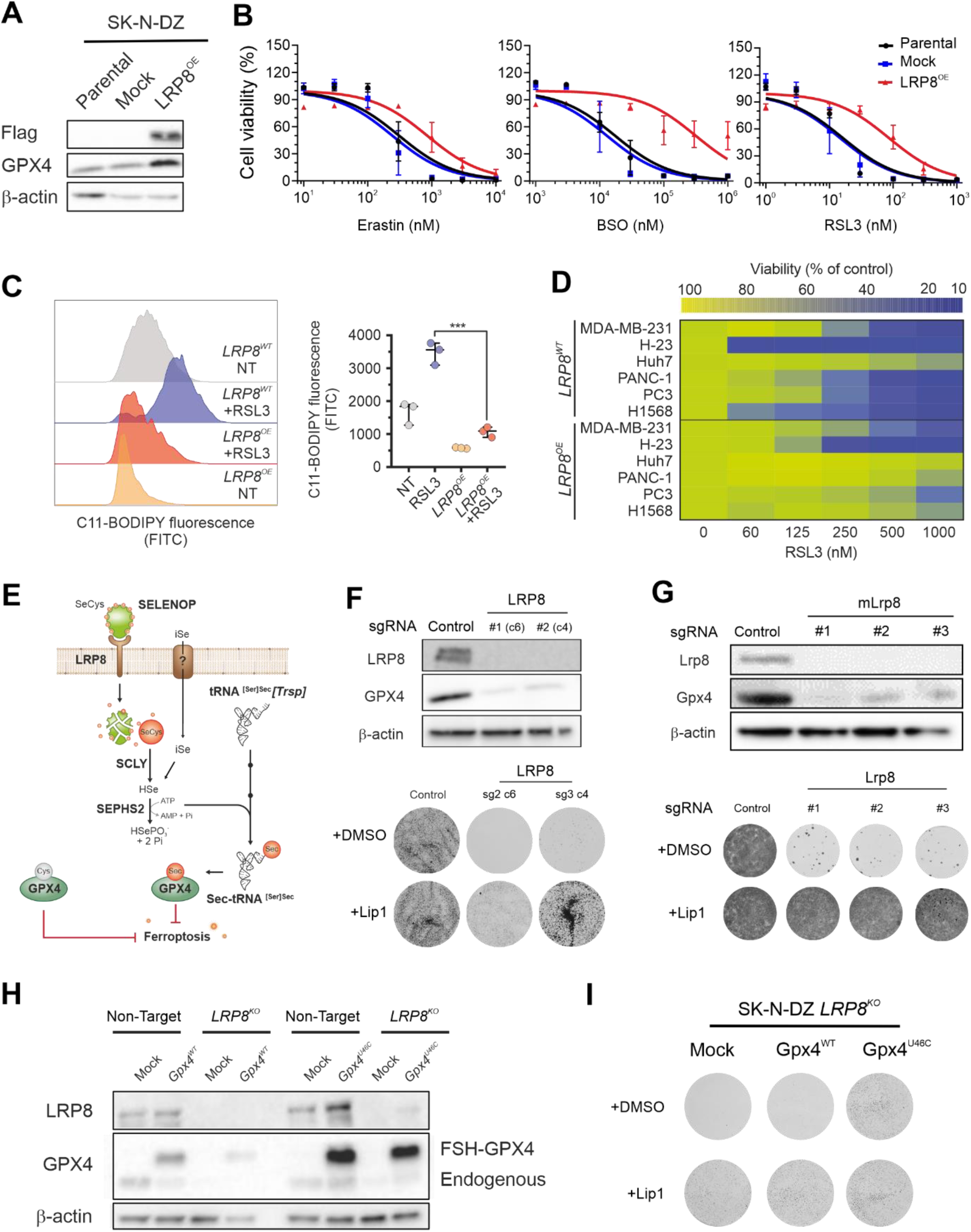
LRP8 loss triggers ferroptosis in *MYCN*-amplified neuroblastoma. **(A)** Immunoblot analysis of FLAG and GPX4 in SK-N-DZ cells overexpressing an empty vector or *hLRP8*-Flag. **(B)** Dose-dependent toxicity of the ferroptosis inducers Erastin, BSO and RSL3 in SK-N-DZ cell lines stably transduced with a vector expressing *hLRP8*-Flag. Data are the mean ± s.e.m. of n = 3 wells of a 96-well plate from three independent experiments. **(C)** Flow cytometry analysis of BODIPY 581/591 C11 oxidation in SK-N-DZ overexpressing *hLRP8*-Flag induced by RSL3 treatment (100 nM, 6 h). **(D)** Heat map depicting the dose-dependent response of RSL3 in a panel of cell lines overexpressing *hLRP8*-Flag. **(E)** Schematic representation of selenium uptake mechanisms. **(F)** Generation and characterization of *LRP8* knockout cell lines using two independent gRNAs. Upper panel, immunoblot analysis of LRP8 expression in cells transduced with gRNA targeting *LRP8*. Lower panel, clonogenic capacity of clonal cell lines either wild-type or knockout for *LRP8* in the presence of Lip-1 (500 nM). **(G)** Recapitulation of LRP8 dependency in a murine model of *MYCN*-amplification. Upper panel depicts immunoblot of Lrp8 and Gpx4 in cells expressing three independent gRNAs targeting *Lrp8*. Lower panel shows the clonogenic capacity of Lrp8-deficient cells and the protective effect of Lip-1 (500 nM). **(H)** Immunoblot analysis of FLAG and GPX4 in SK-N-DZ cells overexpressing flag-tagged WT or an U46C variant of GPX4 in a wild-type and knockout *LRP8* background. **(I)** Clonogenic capacity of SK-N-DZ *LRP8* knockout cells expressing an empty vector, WT or an U46C variant of GPX4. Clonogenic assays depicted are representative of three independently performed experiments with similar results.

Next, we set out to explore the mechanism by which LRP8 protects cells from ferroptosis. Previous studies have established that members of low-density lipoprotein (LDL) receptor-related protein, including LRP2 and LRP8, are receptors for the selenium carrier protein SELENOP (*21, 22*), indicating that modulation of selenocysteine metabolism could be the major mechanisms by which LRP8 protects cells from ferroptosis (**Fig. 2E**). This hypothesis is in agreement with the finding that overexpression of *LRP8* leads to upregulation of GPX4 at the protein level without impacting RNA levels, pointing towards an underlying post-transcriptional regulation (**Fig. 2A** and fig. S1F and 2F). Next, to address the requirement of LRP8, we generated LRP8-deficient SK-N-DZ cells and observed that these cells, without additional stressors, readily underwent massive ferroptosis-like cell death in the absence of ferroptosis-inhibiting compounds (**Fig. 2F**). To further corroborate this observation and provide mechanistic support for targeting LRP8 in neuroblastoma, we deleted *Lrp8* in the murine TH-MYCN neuroblastoma mouse model (**Fig. 2G**) and in a larger panel of neuroblastoma cell lines with and without *MYCN* amplification (fig. S2G-I). These additional models fully recapitulated the observation that LRP8 is essential to support the viability not only of human neuroblastoma cell lines but that it is also essential in a well-defined neuroblastoma genetic model driven by MYCN (fig. S2H-I).

Given the important role of selenocysteine metabolism in preventing ferroptosis, we next asked if any correlation with the clinical outcome can be observed in cohorts of pediatric neuroblastoma patients. First, we showed that high *GPX4* expression is a robust predictor of poor survival, arguing that ferroptosis could indeed be a major tumor-suppressing mechanism in neuroblastoma (fig. S3). Although *LRP8* expression itself was not associated with event-free survival, expression of several members of the selenium/selenocysteine metabolic pathway downstream of LRP8 as well as LRP8 codependent enzymes (i.e, *SEPHS2, PSTK, EEFSEC, SECISBP2*) showed strong correlation with poor disease outcome, a finding that is in line with the generally high expression of members of the selenocysteine biosynthetic pathway in high-risk neuroblastoma (fig. S3). These data suggest that the efficient flux of selenium/selenocysteine through this pathway is a strong determinant for poor survival of neuroblastoma patients and may mediate therapy resistance. This hypothesis also agrees with our observation that SK-N-DZ cells were unable to grow in selected batches of fetal bovine serum (FBS) unless supplemented with exogenous sources of selenium or ferroptosis-inhibiting compounds (fig. S4A). Strikingly, we could identify that FBS unable to support the growth of this cell line was not devoid of total selenium but showed remarkably low levels of SELENOP (fig. S4B). Accordingly, the protective effect conferred by the overexpression of *LRP8* was dramatically reduced in such conditions (fig. S4C). Unequivocal evidence for a direct link between LRP8, selenocysteine, GPX4 and cell survival was provided by showing that overexpression of wild-type GPX4 was unable to rescue cell viability upon loss of *LRP8*, but that expression of GPX4 (U46C), a mutant that retains catalytic function but bypasses the requirement of selenium for efficient translation (*23*), could fully restore cell viability (**Fig. 2H, I**). In line with this, knockdown of selenophosphate synthetase 2 (*SEPHS2*), a central enzyme in the metabolization of selenium, significantly reduced ferroptosis resistance of cells overexpressing *LRP8* (fig. S4D). Additional proof that ferroptosis is induced in LRP8-deficient cells was provided by demonstrating that lipid peroxidation can be triggered in the absence of LRP8 and that this is prevented by the addition of ferrostatin-1 (Fer1) or by the overexpression of the GPX4-independent ferroptosis suppressor *AIFM2* (*FSP1*) (fig. S4E, F) (*15*). Taken together, our results suggest that targeting SELENOP uptake via LRP8 could be a valuable strategy for selectively triggering ferroptosis in *MYCN*-amplified neuroblastoma.

### Characterization of selenium uptake mechanisms

The observation that *MYCN*-amplified neuroblastoma cells are highly dependent on LRP8 for selenium uptake is unanticipated, given that alternative forms and uptake mechanisms exist (*24*). Moreover, the speciation analysis of FBS showed that selenite and selenate were present, both at approximately 1-2 nM, but these species were unable to sufficiently support selenocysteine biosynthesis and consequently unable to suppress ferroptosis in neuroblastoma (fig. S4B). Our *in vitro* rescue experiments suggested that additional 20-fold excess of selenite was required to rescue viability (fig. S4B). These findings led us to hypothesize that alternative mechanisms of selenium provision must be inefficient in neuroblastoma cells. In order to gain insights into these alternative mechanisms, we devised two CRISPR-based screens focusing on solute carriers (SLCs), as these transporters are the major mediators of soluble metabolite uptake into cells (*25*). First, we took advantage of the fact that SK-N-DZ requires supplementation of selenium for proliferation in SELENOP poor conditions. Therefore, we supplemented specific compounds able to support the growth of this cell line, namely selenocystine, selenite, beta-mercaptoethanol and selenomethionine, and compared gRNA representation under these conditions (**Fig. 3A**). To allow such an assessment, we normalized gRNA representation of each condition to that of the selenomethionine condition, reasoning that the complete absence of selenium could impair proliferation and thus hinder the identification of relevant transporters. Analysis of the selenocystine and selenite supplementation conditions revealed that gRNAs targeting *SLC7A11* and *SLC3A2* were the most strongly depleted. This indicates that system Xc-, in addition to its previously reported role in indirect uptake (*26, 27*), is a major contributing factor for the direct uptake of inorganic and organic forms of selenium (**Fig. 3B**). Additionally, our screen also revealed that the sulfate transporter SLC26A6 contributes to the uptake of selenite (**Fig. 3B**).

**Fig. 3.**
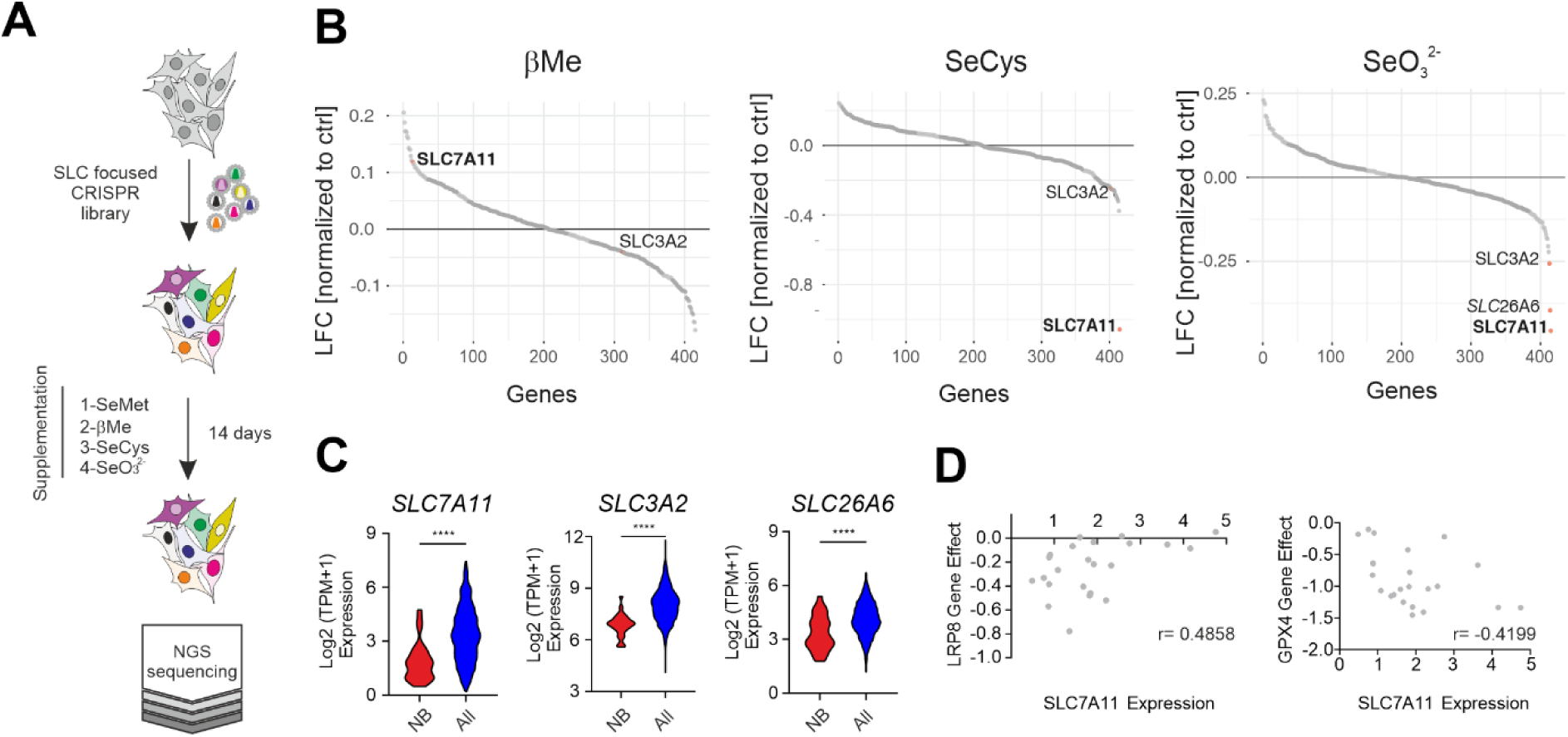
SLC-focused CRISPR screen identifies alternative selenium uptake mechanisms. **(A)** Schematic representation of the SLC-focused CRISPR knockout screen in the SK-N-DZ cell line grown in the presence of defined selenium sources. **(B)** CRISPR gene log2 fold change in SK-N-DZ cells grown in defined supplemented media. **(C)** Comparison of *SLC7A11* expression in a panel of 23 neuroblastoma cell lines against 1349 non-neuroblastoma cell lines demonstrating the lineage-specific lower expression of *SLC7A11, SLC3A2* and SLC26A6 (www.depmap.org). **(D)** Dot plot depicting the correlation of the dependency of neuroblastoma cell lines on LRP8 and GPX4 (CERES score of −1 means full dependency based on CRISPR–Cas9 knockout screening data) and the expression levels of *SLC7A11* in a panel of 27 neuroblastoma cell lines (depmap portal; https://depmap.org/portal/). Cell lines with low expression of *SLC7A11* were found to be dependent on LRP8 (Pearson correlation *r:*0.4858).

In an alternative screen aiming to identify SLC synthetic lethal interactions with LRP8, we used cells in which the deletion of *LRP8* did not lead to the loss of viability (HT1080 and A375) (fig. S5A-D). Despite being viable upon genetic loss of *LRP8*, these cell lines still display a high sensitivity towards several ferroptosis inducers (fig. S5B, D). Similarly, screens in both cell lines retrieved the system Xc-subunits SLC7A11 and SLC3A2 as synthetic lethality with LRP8 deficiency (fig. S5E). Surprisingly, the SLC26A6 transporter was not identified in these settings, suggesting that this transporter might be only relevant in specific cell types, indicating redundancy with other members of the SLC26A family. In line with these observations, neuroblastoma cell lines show overall reduced expression levels of both the system Xc-(*SLC7A11* and *SLC3A2*) and *SLC26A6*, compared to other cancer cell lines (**Fig. 3C**). Furthermore, analysis of publicly available data (depmap portal) showed that LRP8 dependency presents a significantly positive correlation with *SLC7A11* expression in neuroblastomas, whereas, curiously, GPX4 dependency presented an inverse correlation (**Fig. 3D**). Such correlations were not obvious for SLC26A6. Altogether, this data suggests that the low capacity for the uptake of inorganic selenium is the likely cause for the increased dependency on LRP8 in neuroblastoma.

### SLC7A11 activity is toxic to MYCN-amplified neuroblastoma

Next, we decided to further investigate the role of system Xc-, reasoned by its general importance in multiple cellular contexts. We rationalized that overexpression of *SLC7A11* in LRP8-dependent cell lines could be sufficient to rescue them from cell death, as this should allow them to exploit alternative selenium sources. As initially foreseen, overexpression of

*SLC7A11* led to a marked increase in GPX4 protein level, an effect phenocopied by the thiol donor βMe (Fig. 4A). Even though GPX4 was robustly upregulated, we, surprisingly, observed that *SLC7A11* overexpression led to a profound growth defect in neuroblastoma cells. This effect was specific to SLC7A11 activity, as it was fully rescued by the addition of the system Xc-inhibitor Erastin (**Fig. 4A**). Metabolomics and RNA-seq analysis revealed that increasing SLC7A11 activity resulted in a marked upregulation of intracellular thiols and was accompanied by the induction of several oxidative stress-related genes that are controlled by ATF4 and NRF2 (**Fig. 4B, C and** fig. S6B, C). To provide additional insight into potential causes of the SLC7A11-mediated toxicity in neuroblastoma, we generated SK-N-DZ cells carrying a Dox-inducible *SLC7A11* vector. This model system showed that upon doxycycline treatment *SLC7A11*-expressing cells lose proliferative capacity and eventually die (fig. S6A). These results are in agreement with reports demonstrating the toxic impact of cysteine overload on mitochondrial function and the induction of increased reductive stress (*28, 29*). This notion was further corroborated by our analysis of mitochondrial function (**Fig. 4D, E**) showing that oxygen consumption was significantly impaired in cells overexpressing *SLC7A11*. This was also mirrored by the loss of intermediates of the tricarboxylic acid (TCA) cycle and by a noticeable depletion of nucleotide triphosphates (NTP) (**Fig. 4B, C**). Curiously, in contrast to the metabolic profile of *SLC7A11*-overexpressing cells, cells overexpressing *LRP8* presented an inverse response. Specifically, *LRP8*-overexpressing cells showed an increase in NTP levels, most prominent for ATP and UTP, as well as higher levels of NADP and GSH, indicating an increased cellular reducing capacity (fig. S6D-I). These data also suggest a potential role for selenocysteine metabolism in suppressing system Xc-activity. Currently, it is not clear how the expression of LRP8 could regulate system Xc-activity, but, based on our scRNA-seq data, we can hypothesize that this regulation could be mediated post-transcriptionally by the downregulation of an RNA-binding protein, *YBX3*, that was previously reported to bind to the mRNAs of both *SLC7A11* and *SLC3A2* and to stabilize *SLC3A2* (*30*). This hypothesis is supported by the observed regulation of *YBX3* in several CRISPRa clusters (fig. S1E, H).

**Fig. 4.**
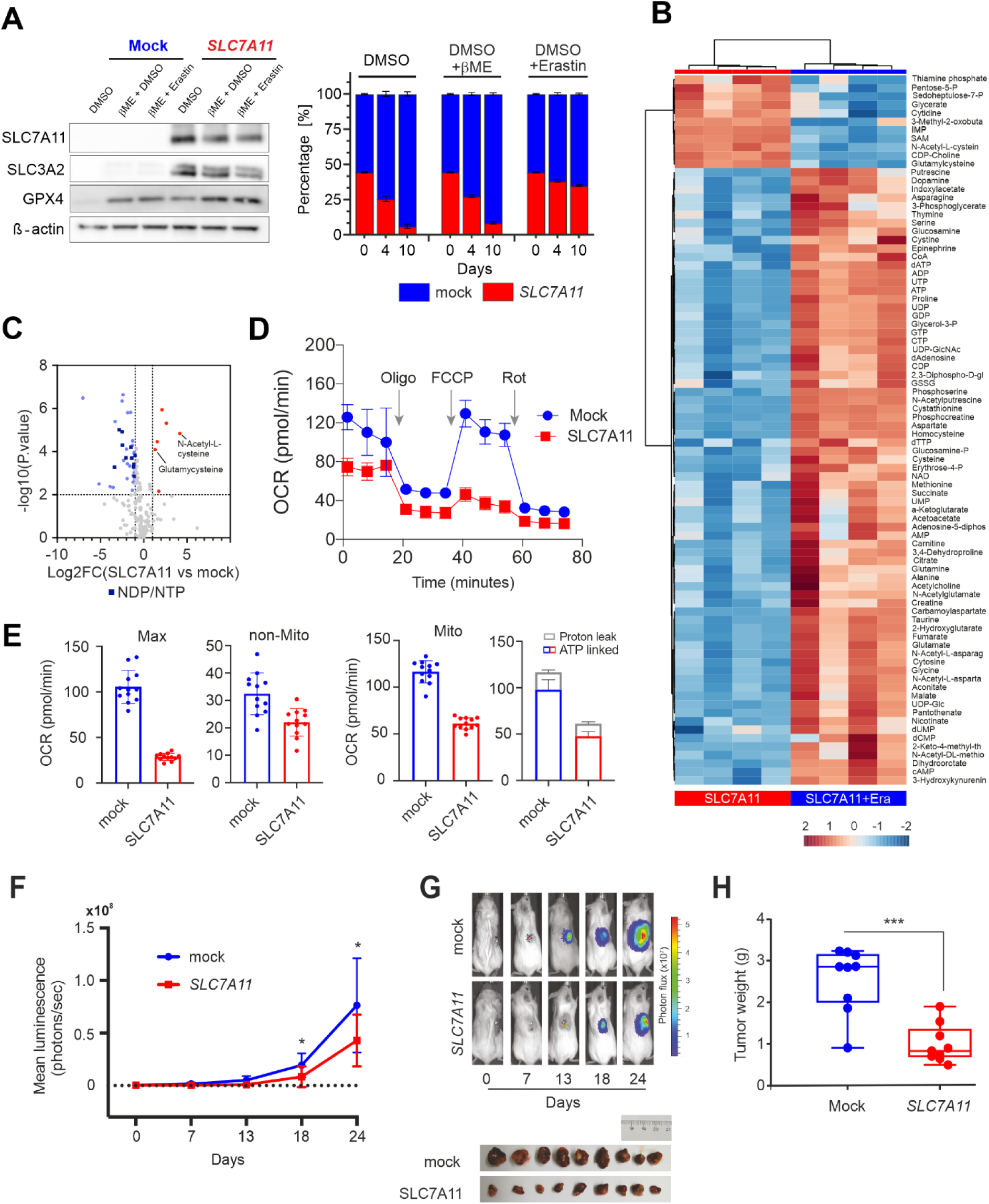
SLC7A11 activation drives metabolic collapse in *MYCN*-amplified neuroblastoma. **(A)** Generation and characterization of cells overexpressing *SLC7A11*. Immunoblot analysis of SLC7A11 and SLC3A2 from SK-N-DZ cells overexpressing *SLC7A11* or mock. Cell competition assay of SK-N-DZ cell line overexpressing *SLC7A11* or mock controls. For the experiment, cells were seeded at a ratio of 50/50 and 50,000 events were measured via flow cytometry at the depicted time points. Rescue experiments were performed in the presence of 50 μM beta-Mercaptoethanol (βME) and 0.5 μM Erastin. Bars display percentage of eGFP-(blue) and Scarlet-cells (red) with n=1, or means +/-s.e.m with n=3. **(B)** Heatmap showing 84 metabolites that are significantly different between *SLC7A11*-overexpressing SK-N-DZ cells with and without Erastin (0.5 μM). Abundance is represented as log2-transformed normalized peak intensity relative to row mean. **(C)** Volcano plot showing log2 fold change of metabolites in *SLC7A11*-overexpressing SK-N-DZ cells treated with or without Erastin (0.5 μM). Metabolites that are significantly different (p≤0.05) with log2 fold change ≥1 or ≤-1 are highlighted. NDP/NTP are marked in dark blue. **(D)** Mitochondrial Stress Assay of SK-N-DZ cells expressing empty vector or *SLC7A11*. **(E)** Mitochondrial parameters derived from for assay shown in (**D**). **(F)** Tumor growth of orthotopically implanted SK-N-DZ cells overexpressing *SLC7A11* (red, *n* = 9) or mock controls (blue, *n* = 9). Values are mean ± s.e.m; statistical comparisons were done using t-tests. * P < 0.05. **(G)** Representative luminescence images and photographs of tumors from each group from (**F**) at endpoint. **(H)** Tumor weight of orthotopically implanted SK-N-DZ cells overexpressing *SLC7A11* (red, *n* = 9) or mock controls (blue, *n* = 9).

Moreover, we could demonstrate that the toxic effect of high *SLC7A11* expression is not limited to the SK-N-DZ cell line, as it could also be recapitulated in the murine cell line derived from the TH-MYCN model. Interestingly, cell lines that do not carry the *MYCN* amplification (SK-N-FI and SH-SY5Y) were mostly refractory to the growth inhibitory effect of *SLC7A11* overexpression (fig. S7A, B). Additional support was provided by using the SH-EP (MYCN-ERT2) model, where we could demonstrate that decreased cell fitness already reported by MYCN activation (*31*) is aggravated by *SLC7A11* overexpression and rescued by Erastin treatment (fig. S7C, D). Finally, growth impairment upon *SLC7A11* overexpression was not restricted to cell lines grown *in vitro* as orthotopic *in vivo* models of neuroblastoma also exhibited a marked decrease in tumor growth upon *SLC7A11* induction (**Fig. 4F-H**). Together, our results suggest that the LRP8 dependency could emerge during tumor evolution as a result of a potential growth advantage of neuroblastoma with low activity of system Xc^-^.

### LRP8 is required for the initiation and maintenance of neuroblastoma in vivo

Given the important role played by the selenocysteine metabolism in preventing ferroptosis in highly aggressive neuroblastoma subtypes with *MYCN* amplifications, we asked if upstream targeting of this pathway could afford a detrimental effect on neuroblastoma growth *in vivo*. We took advantage of our orthotopic animal model in which SK-N-DZ cells were implanted in the adrenal gland of NOD.Cg-Prkdc^scid^Il2rgtm1^Wjl^/SzJ (NSG) mice (*12*). Briefly, cells were grown in the presence of liproxstatin-1 (Lip-1) before implantation to prevent ferroptotic cell death of LRP8-deficient and -proficient neuroblastoma cells. Subsequently, SK-N-DZ cells were implanted and tumor growth was monitored using *in vivo* bioluminescence imaging (**Fig. 5A**). LRP8 deficient tumors showed a marked decrease in tumor growth and, compared to LRP8 proficient cells, showed a significantly increased overall survival of mice (76d +/-vs. 31d +/-) with a median survival of 29d vs. 40days (**Fig. 5B-D**). Next, we explored the therapeutic potential of targeting LRP8 in established SK-N-DZ neuroblastoma. For this, LRP8 proficient (LRP8^WT^) and deficient (LRP8^KO^) tumors were both allowed to grow in the presence of in vivo active Lip-1 (**Fig. 5E**) with the former ones serving as controls (**Fig. 5E**, top, red). After LRP8-deficient neuroblastomas were established (seven days post-implantation with five days Lip-1 treatment), mice were randomized into two groups. Lip-1 treatment was continued in the yellow group, while in the blue group, the Lip-1 injections and thus ferroptosis inhibition were stopped (**Fig. 5E**, bottom). Analysis of the three cohorts after 14 days post-implantation showed that both LRP8 deficient xenograft groups (blue, yellow) showed impaired tumor growth compared to WT controls (red) despite Lip-1 mediated ferroptosis inhibition (**Fig. 5F, G**). Importantly, Lip-1 withdrawal in the randomized LRP8^KO^ group significantly further reduced neuroblastoma growth in the established tumor setting (**Fig. 5F,G**). The data above support the requirement of LRP8 for the establishment and maintenance of MYCN amplified SK-N-DZ neuroblastoma by restricting ferroptosis. This is further confirmed by the fact that the LRP8^KO^ tumors show robust loss of GPX4 protein (**Fig. 5H**). Collectively, our data indicate that LRP8 is required to prevent ferroptosis by maintaining high levels of GPX4 in *MYCN*-amplified orthotopic neuroblastoma models and suggest that inhibiting the SELENOP/LRP8 axis could be a novel and selective strategy to trigger ferroptosis and thereby limit tumor growth in neuroblastoma cells (Fig. 5I).

**Fig. 5.**
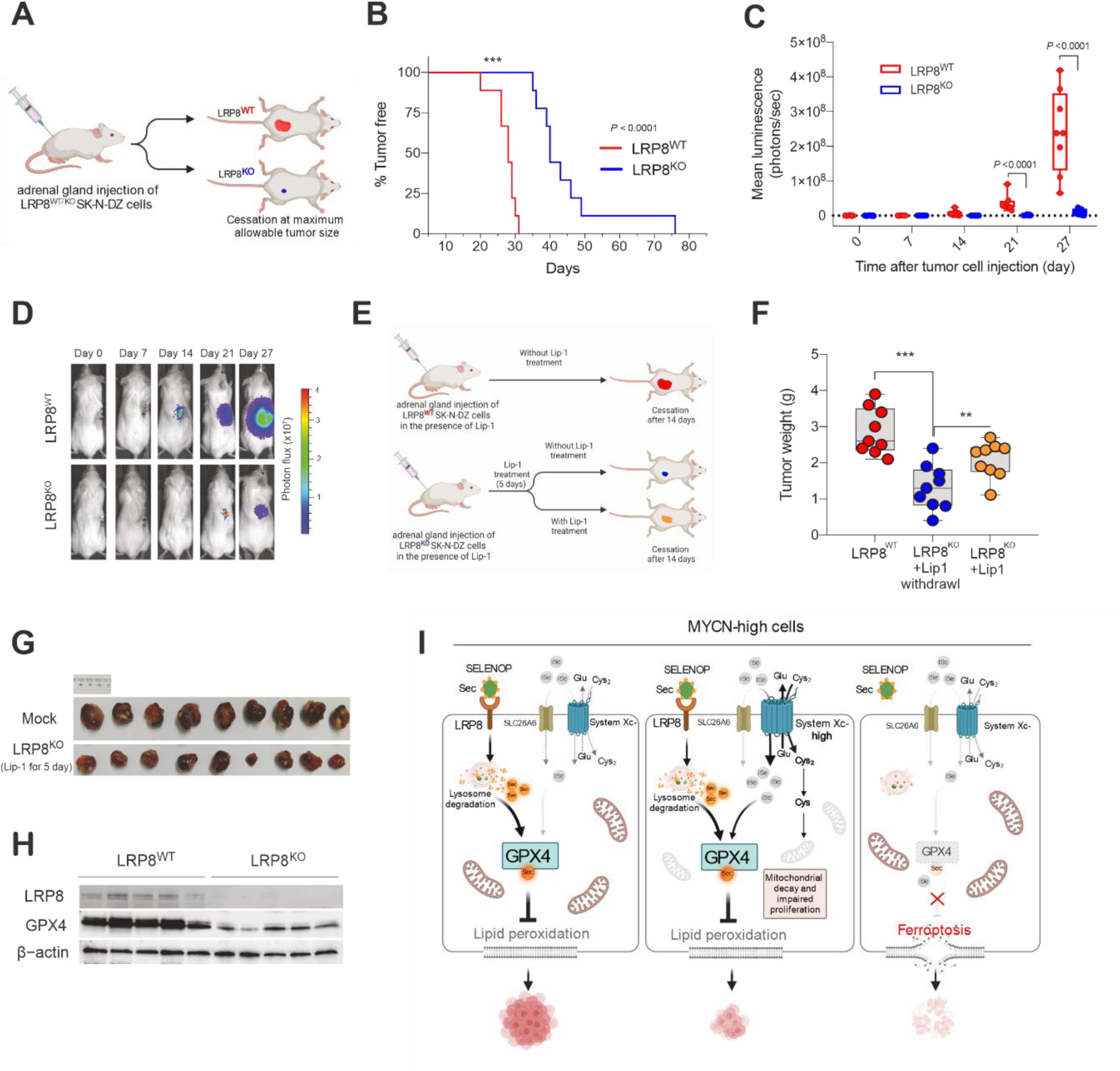
LRP8 is essential for orthotopic neuroblastoma growth. **(A)** Schematic representation of the orthotopic implantation of control (LRP8^WT^) or LRP8-deficient (LRP8^KO^) SK-N-DZ cell lines. **(B)** Kaplan-Meier plot displaying tumor-free survival (TFS) for mice injected orthotopically with LRP8^WT^ (blue, *n* = 9) or LRP8^KO^ (red, *n* = 9) SK-N-DZ cells. t-test was conducted for statistical analysis. **** P < 0.0001. **(C)** Tumor growth upon orthotopic implantation of LRP8^WT^ (blue, *n* = 9) or LRP8^KO^ (red, *n* = 9) of SK-N-DZ cell line. Values are mean with SEM; t-test was conducted for statistical analysis. **** P < 0.0001. **(D)** Representative luminescence images from each group sown in (c). **(E)** Outline of the orthotopic implantation and treatment scheme with Liproxstatin-1 (Lip-1) of LRP8^WT^ and LRP8^KO^ SK-N-DZ cells **(F)** Tumor weight of orthotopically implanted LRP8^WT^ (blue, *n* = 9) or LRP8^KO^ (red and gray, *n* = 9 each) SK-N-DZ cells. All mice were treated with Lip-1 for 5 days. After this, treatment was ceased for LRP8^WT^ (blue) and LRP8^KO^ cohorts (red) or maintained for an additional 14 days (yellow), see methods for details. Tumors were analyzed at the end point. Values represent mean with s.e.m; t-test was conducted for statistical analysis. **(G)**. Representative images of tumors in the control (blue) and LRP8^KO^ groups (yellow) treated with Lip-1 for 5 days after implantation. **(H)** Immunoblot analysis of LRP8 and GPX4 levels from orthotopic tumors of LRP8^WT^ or LRP8^KO^ SK-N-DZ cells, treated with Lip-1 for 5 days after implantation followed by 14 days after randomization. **(I)** Schematic representation for the proposed model of LRP8 inhibition essentiality. Comparison of selenium /selenocysteine uptake mechanisms in proliferating MYCN-amplified cells, depicting the contribution of primarily LRP8/SELENOP supporting selenoprotein translation (left panel). Activation of SLC7A11 mediated uptake of selenium/selenocysteine in *MYCN*-amplified neuroblastoma leads to progressive mitochondrial decline and impaired proliferation (central panel). Inhibition of LRP8 in proliferative and SLC7A11/SLC26 low conditions selectively trigger ferroptosis in *MYCN*-amplified neuroblastoma (right panel). SELENOP, Selenoprotein P; iSe, inorganic selenium; SeCys, Selenocysteine.

## Discussion

The present study demonstrates that blocking selenium/selenocysteine uptake mechanisms could be an attractive strategy to disrupt GPX4 function specifically and selectively induce ferroptotic cell death in *MYCN*-amplified neuroblastoma. We demonstrate here that selenium/selenocysteine can be obtained by cancer cells via multiple routes: one dependent on the SELENOP/LRP8 axis and alternatively via the activity of cyst(e)ine/sulfate transport systems such as system Xc-(SLC7A11/SLC3A2) and members of the SLC26A family. Characterization of these two pathways in neuroblastoma uncovered that they pose a fundamentally distinct metabolic burden for the *MYCN*-amplified subgroup, where activation of system Xc-leads to mitochondrial functional decline and impaired proliferation favouring the development of cancer cells with low system Xc-activity. The tradeoff for this aggressive and fast proliferative state generates an unexpected dependency on LRP8 and thus unravels a rational strategy to selectively disrupt GPX4 and elicit ferroptosis in *MYCN*-amplified entities. These recognitions could have broader implications, as a recently published cancer dependency map of pediatric tumors identified *LRP8* as an essential gene in pediatric Ewing sarcoma and medulloblastoma, all entities associated with *MYCN* amplifications (*32*). Moreover, given the largely unsuccessful repurposing of adult oncology drugs for the treatment of neuroblastoma and other childhood malignancies, these discoveries pave the way for exciting translational opportunities. Thus, targeting of LRP8, for example, by the development of LRP8/SELENOP neutralizing antibodies, may overcome the limitations of strategies that aim to directly inhibit GPX4 or other members of the selenocysteine metabolic network, which could be limited by extensive organ toxicity as demonstrated by mouse genetic studies (*33*). Additionally, given the recognition that GPX4 inhibition can also impair T-cell function (*34*), but also contribute to the CD8+ T-cell mediated cancer cell killing (*35, 36*), it is important to consider that immune cells, like CD8+ T-cells lack *LRP8* expression. Thus, targeting LRP8 could critically impair GPX4 function in MYCN tumors or metastasis while sparing immune cells and could therefore be seen as a potent combinatorial strategy with immunotherapies.

## Funding

J.P.F.A. acknowledges the support of the Junior Group Leader program of the Rudolf Virchow Center, University of Würzburg, the Deutsche Forschungsgemeinschaft (DFG), FR 3746/3-1, SPP2306 (FR 3746/6-1) and the CRC205 (INST 269/886-1). This work was also supported by the SPP2306 (A.T. and H.A.), FOR2674 and SFB873 funded by the Deutsche Forschungsgemeinschaft (A.T.); the DKTK joint funding project “RiskY-AML”; the “Integrate-TN” Consortium funded by the Deutsche Krebshilfe, the Dietmar Hopp Foundation (A.T.). M.C. is supported by the DFG CO 291/7-1 and the SPP2306 (CO 297/9-1 and CO 297/10-1) as well as the European Research Council (ERC) under the European Union’s Horizon 2020 research and innovation programme (grant agreement No. GA 884754). J.Z. is supported by a Humboldt Postdoctoral Fellowship. A.S., L.S. and F.C.E.V. are supported by SPP2306 (SCHU 2670/4-1) and FOR2314 (SCHU 2670/2-2). MF and CB were supported by the Deutsche Forschungsgemeinschaft (DFG; grant no. BA 6984/1-1 and grant no. FI 1926/2-1 to M.F., and as part of the SFB 1399 to M.F.), the Förderverein für krebskranke Kinder e.V. Köln (endowed chair to M.F.), and the German Ministry of Science and Education (BMBF) as part of the e:Med initiative (grant no. 01ZX1603 and 01ZX1607 to M.F.). G.S.F acknowledges funding from the Austrian Academy of Sciences. E.G was supported by a Marie Sklodowska-Curie fellowship (MSCA-IF-2014-661491). U.Y. and H.A. thank the DKFZ Single-Cell Open Lab (scOpenLab) for assistance with the single-cell CRISPR-activation screen. Illustrations in Figures 1B, E, and 5A, E, I were created with BioRender.com.

## Author Contributions

J.P.F.A., A.T. and H.A. conceived the project. J.P.F.A., H.A. designed experiments and wrote the manuscript. Z.C., H.A., U.Y. performed most experiments described herein and related analyses. J.B., F.P.F., A.F.S., T.N.X.S., J.V., N.A., A.F., P.Z., A.P. supported with the in vitro experiments, generation of cell lines and analyses thereof. U.Y. performed the analyses of the CRISPRa screens and scRNA-seq. H.A., E.E. performed the in vivo experiments and implemented related analyses. C.B. and M.F. generated the neuroblastoma cohort data previously, and performed the analyses depicted here. W.S., F.C.E.V., L.S. and A.S. supported with the preparation, analysis and interpretations of metabolomics data. J.Z., M.C., G.B., M.E., E.G. and G.S.F. generated and provided antibodies, cell lines and key reagents used in this study. B.M performed and analyzed the selenium speciation analysis. J.P.F.A. and A.T. supervised the project. All authors provided intellectual input and critical edits and approved the manuscript.

**Additional Information**, n.a.

**Fig. S1.**
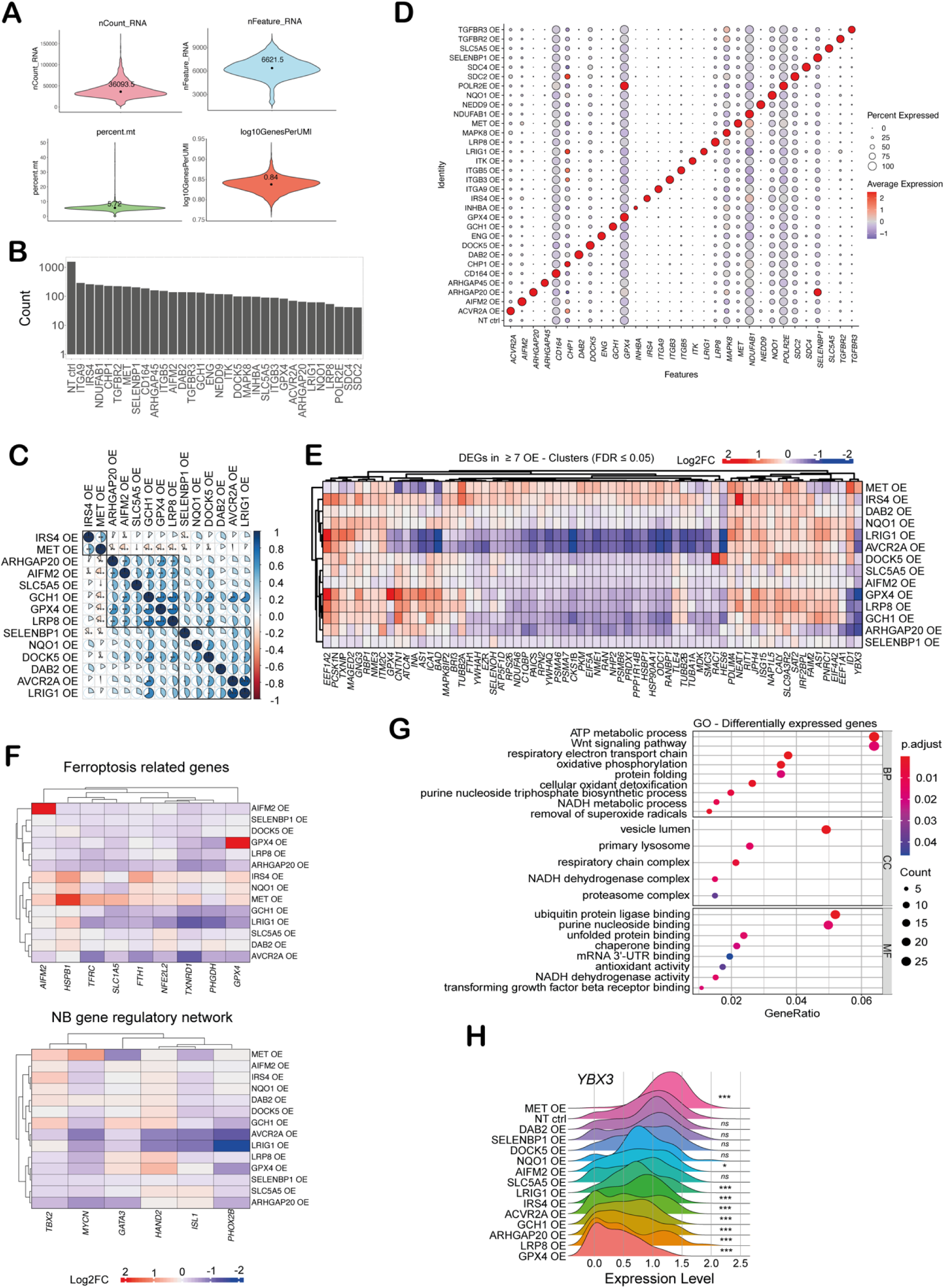
**(A)** Top left: number of unique molecular identifiers (UMIs) per cell in the scCRISPRa sample. Top right: number of detected genes per cell. Bottom left: percentage of mitochondrial reads detected. Bottom right: number of genes per UMI (log-transformed) reflecting library complexity. The median values are printed and highlighted by a dot. **(B)** The number of cells (y-axis, in log-scale) assigned to each scoring hit (indicated on the x-axis) in the single-cell experiment. OE: overexpression, NT ctrl: cells assigned to non-targeting control gRNAs. **(C)** Pairwise comparison of transcriptome similarities of validated CRISPRa clusters and genes derived from the gene expression signature list (based on Spearman correlation). Color intensities and ratios of the pie-chart indicate relative overlap. Rows and columns are hierarchically ordered and related groups are marked with a blue rectangle. \*\*\**P* < 0.001, \*\**P* < 0.01, \**P* < 0.05. **(D)** Dot plot showing expression of target genes (x-axis) in each CRISPRa cluster (y-axis). The size of the dots reflects the fraction of cells within a particular cluster for which the expression of the gene was detected. OE: overexpression. NT ctrl: cluster of cells assigned to non-targeting control gRNAs. **(E)** Changes in gene expression following CRISPRa (clusters represented by each row) of genes that were detected as significantly differentially expressed (FDR ≤ 0.05) in at least seven of the 14 scoring CRISPRa clusters. Columns and rows were hierarchically clustered based on Pearson correlation. **(F)** Changes in expression of ferroptosis related genes (upper) and transcriptional core regulatory circuits maintaining cell state in *MYCN*-amplified neuroblastoma derived from (*37*) (lower) that were detected as significantly differentially expressed in at least one of the CRISPRa clusters represented by each row. **(G)** Gene Ontology (GO) term enrichment analysis conducted on the signature gene list that was derived from the top 50 genes showing the most significant differential expression in each scoring CRISPRa cluster. Selected terms are shown up to FDR ≤ 0.05. BP: biological process, CC: cellular compartment, MF: molecular function. **(H)** Ridge-plot showing the expression of *YBX3* in each scoring CRISPRa cluster. OE: overexpression, NT ctrl: cluster of cells assigned to non-targeting control gRNAs. \*\*\**P* < 0.001, \*\**P* < 0.01, \**P* < 0.05, *ns:* nonsignificant (FDR corrected p-values calculated by Wald tests comparing mean expression values in each CRISPRa cluster versus the NT ctrl cluster).

**Fig. S2.**
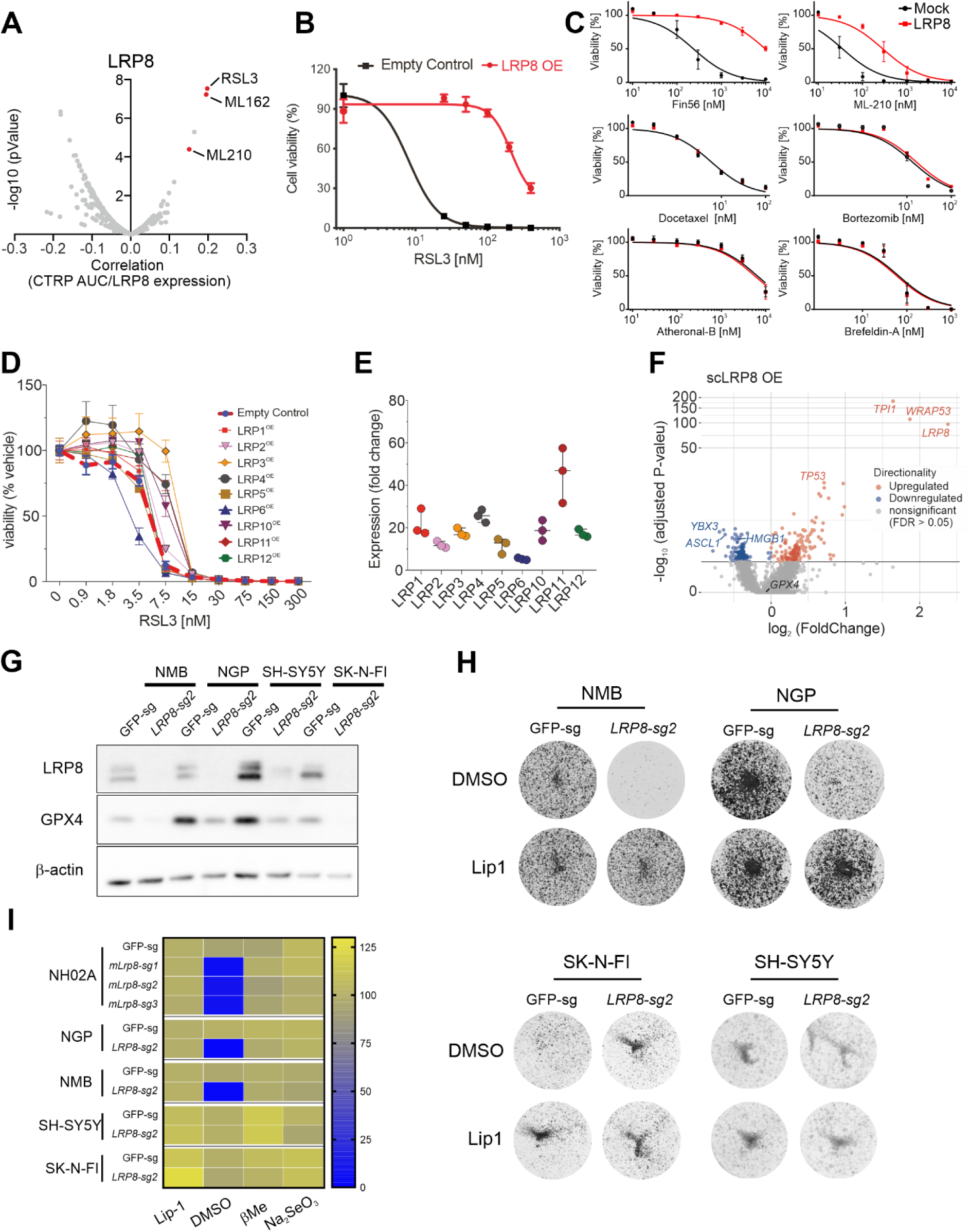
**(A)** High expression of LRP8 is correlated with resistance to GPX4 inhibitors in a panel of 1400 cell lines. Data were extracted from the CTRP/depmap database. **(B)** Dose-dependent toxicity of RSL3 in SK-N-DZ dCas9-SAM cells expressing gRNA targeting the *LRP8* promoter or a control gRNA. **(C)** Dose-dependent response of SK-N-DZ cells overexpressing an empty vector or human *LRP8* (hLRP8) using a panel of cytotoxic drugs. **(D)** dose-dependent toxicity of RSL3 in SK-N-DZ dCas9-SAM cell expressing gRNAs targeting different members of the LRP family. Data are the mean ± s.e.m. of n = 3 wells of a 96-well plate from three independent experiments. **(E)** Expression changes of different LRPs in SK-N-DZ dCas9-SAM cell expressing the respective gRNA. Data are presented as mean ± s.e.m. of n = 3 wells. **(F)** Volcano-plot showing differently expressed genes in the LRP8 overexpression cluster compared to non-targeting control cells. Each dot represents a single gene. Mean expression values (log2 fold-change) are plotted against the significance values (log10 transformed adjusted p-values). Significantly upregulated genes are marked in red while downregulated genes are marked in blue (FDR ≤ 0.05). **(G)** Immunoblot analysis of LRP8 and GPX4 in *MYCN*-amplified (NMB and NGP) and non-amplified (SH-SY5Y and SK-N-FI) neuroblastoma cells transduced with gRNA targeting *Lrp8*. **(H)** Clonogenic capacity of the indicated neuroblastoma cell lines transduced with an gRNA targeting *LRP8* in the presence or absence of the ferroptosis suppressor liproxstatin-1 (Lip-1). **(I)** Heatmap indicating viability of the indicated cell lines transduced with gRNA expression constructs targeting *LRP8* or controls (GFP-sg) in the presence of different ferroptosis inhibitors, including sodium selenite. Data are presented as mean ± s.e.m. of n = 3 wells of a 96-well plate from three independent experiments.

**Fig. S3.**
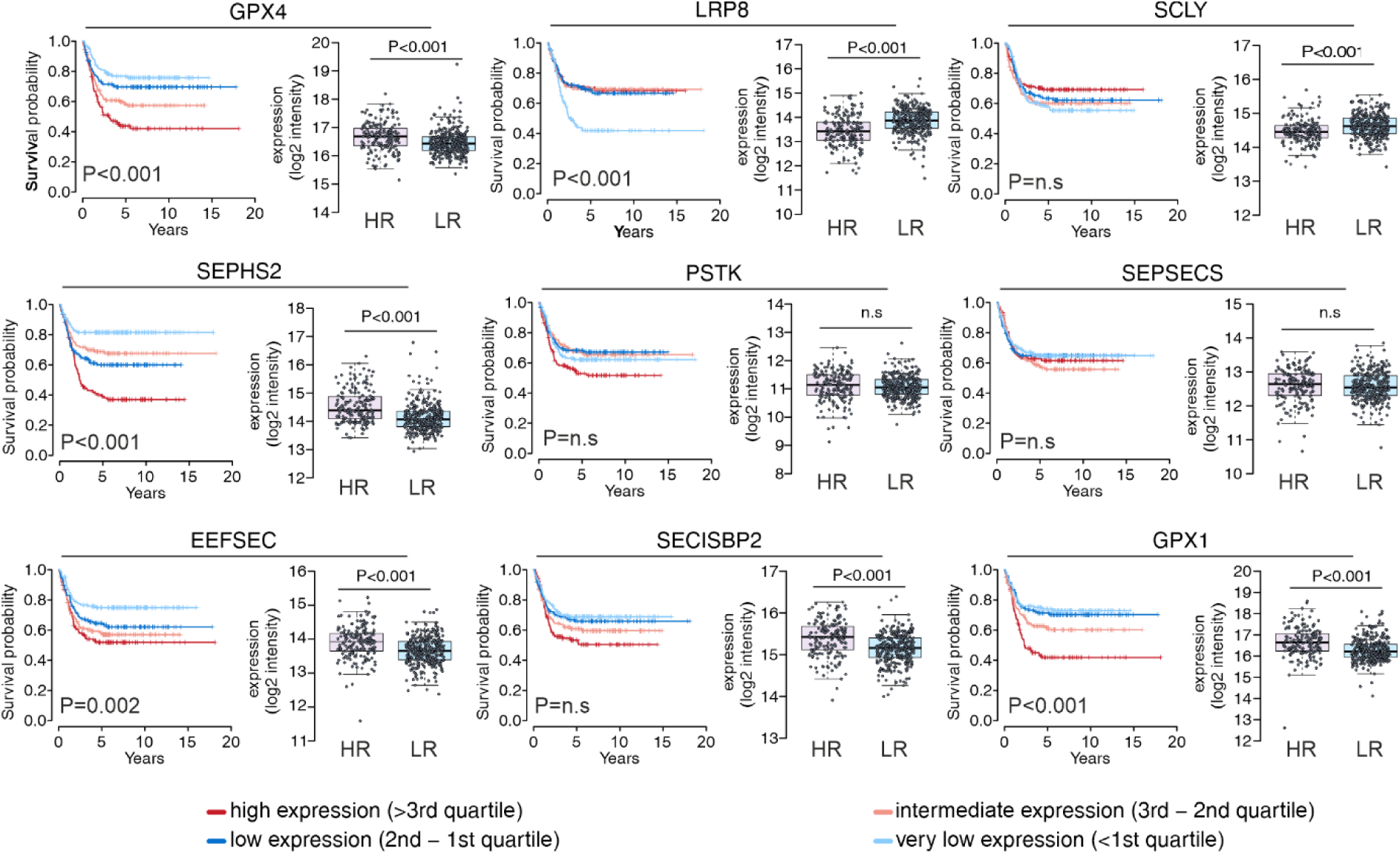
Association between selected genes in the selenium/selenocysteine biosynthetic pathway on Kaplan-Meier survival analysis and expression levels (log2 intensity) in low-risk (LR) and high-risk (HR) neuroblastoma patients (n=459). The p-values were calculated using a two-sided Wilcoxon rank-sum test (boxplots, comparison of gene expression between high and low risk patients) and log-rank test (Kaplan-Meier curves, pairwise comparisons) and Benjamini-Hochberg corrected. All p-values were adjusted for multiple testing (Benjamini-Hochberg).

**Fig. S4.**
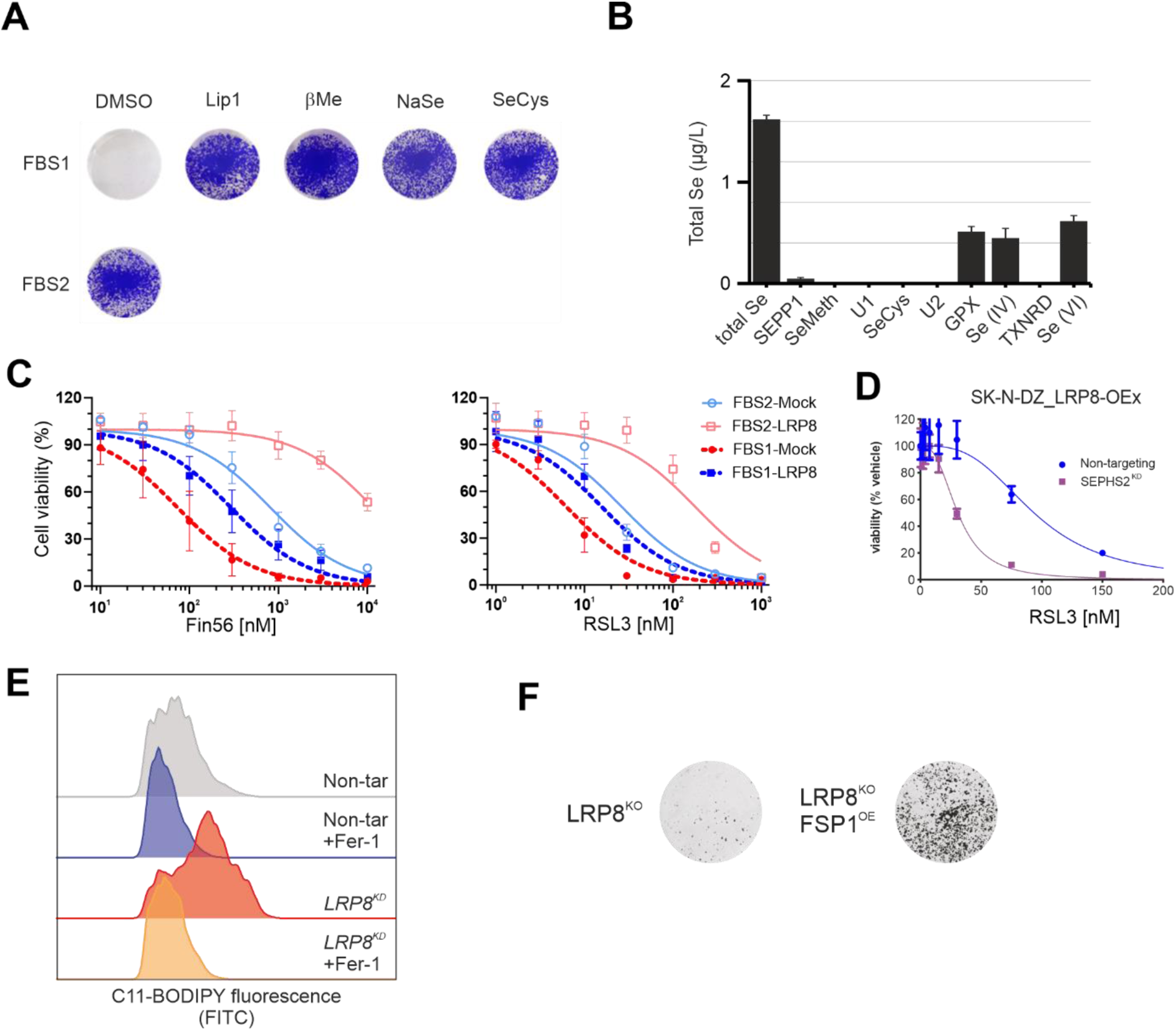
**(A)** Clonogenic capacity of SK-N-DZ cells in two different FBS batches (FBS1 and FBS2). Rescue in FBS1 was carried out using liproxistatin-1 (Lip-1, 500 nM), βMe (50 μM), NaSe (50 nM) and selenocysteine (50 nM). **(B)** Analysis of selenium speciation in FBS lacking growth-supporting capacity of SK-N-DZ cells (FBS1). **(C)** Dose-dependent toxicity of FIN56 and RSL3 in SK-N-DZ expressing a lentiviral construct expressing Flag-*LRP8* (LRP8) or empty vector control (Mock). Viability assays were performed in the two different FBS batches described in (**A**). **(D)** Dose-dependent toxicity of RSL3 in SK-N-DZ cells overexpressing *LRP8* (LRP8-OEx) and co-transfected with siRNA targeting *SEPHS2* (SEPHS2^KD^) or non-targeting control. **(E)** Flow cytometry analysis of C11-BODIPY oxidation in SK-N-DZ *LRP8* knockdown cells (LRP8^KD^) or non-targeting controls in the presence and absence of ferrostatin-1 (1 μM). **(F)** Clonogenic capacity of SK-N-DZ *LRP8* knockout cells (LRP8^KO^) expressing a FSP1 overexpression construct (FSP1^OE^) or empty vector control.

**Fig. S5.**
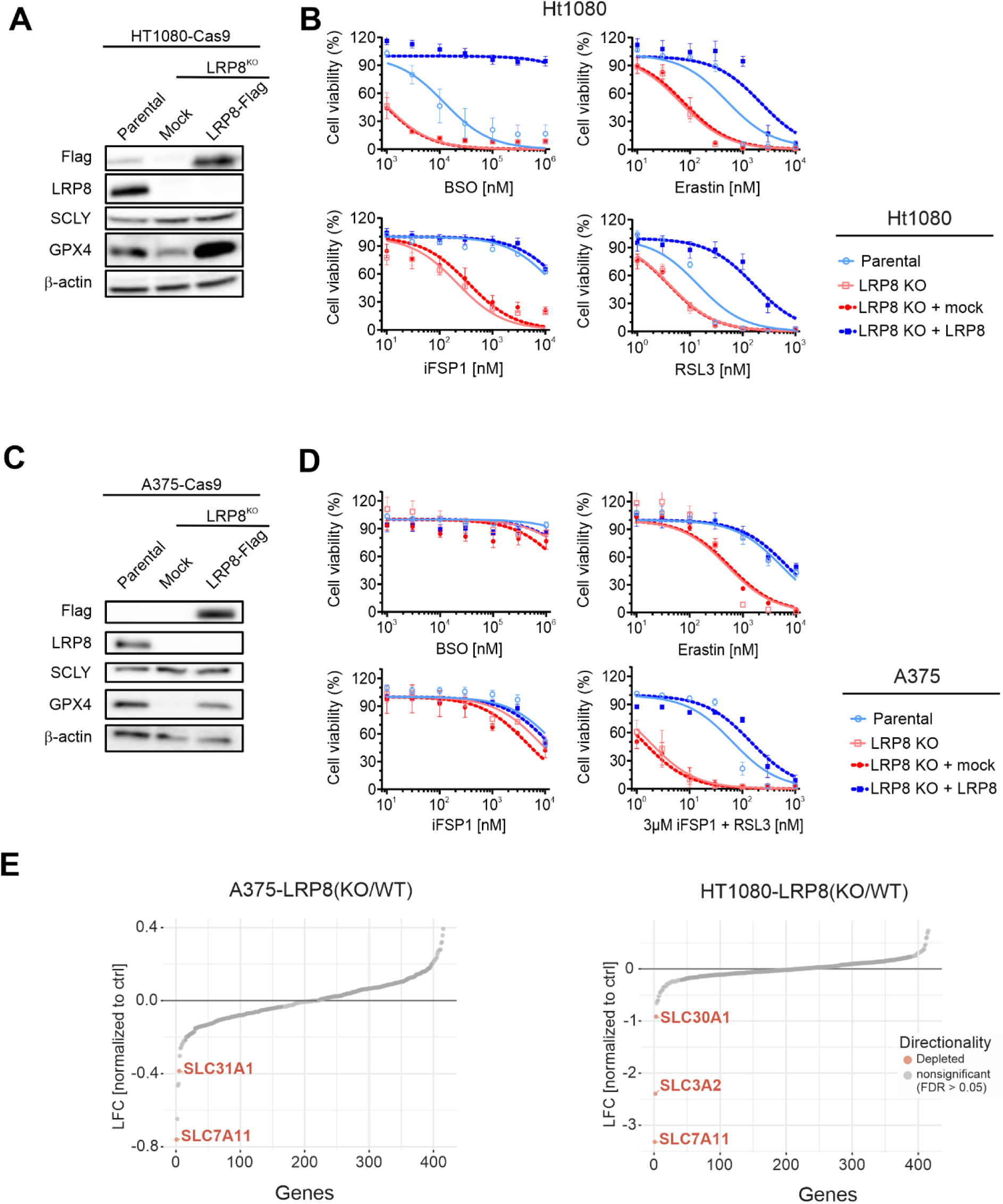
Immunoblot analysis of LRP8, SCLY, GPX4 and Flag in LRP8-deficient HT1080 (**A**) and A375 (**C**) cells overexpressing *hLRP8*-Flag or empty vector (Mock). Parental cell lines only expressing Cas9 are shown as control. Dose-dependent toxicity of the ferroptosis inducers iFSP1, Erastin, BSO and RSL3 in LRP8-deficient (LRP8 KO) HT1080 (**B**) and A375 (**D**) cell lines stably transduced with a vector expressing *hLRP8*-Flag. Parental cell lines only expressing Cas9 are shown as control. Data are the mean ± s.e.m. of n = 3 wells of a 96-well plate from three independent experiments. **(E)** Results of a CRISPR deletion screen conducted in H1080 and A375 cell lines displayed as log2 fold change between LRP8-deficient and wild-type cells.

**Fig. S6.**
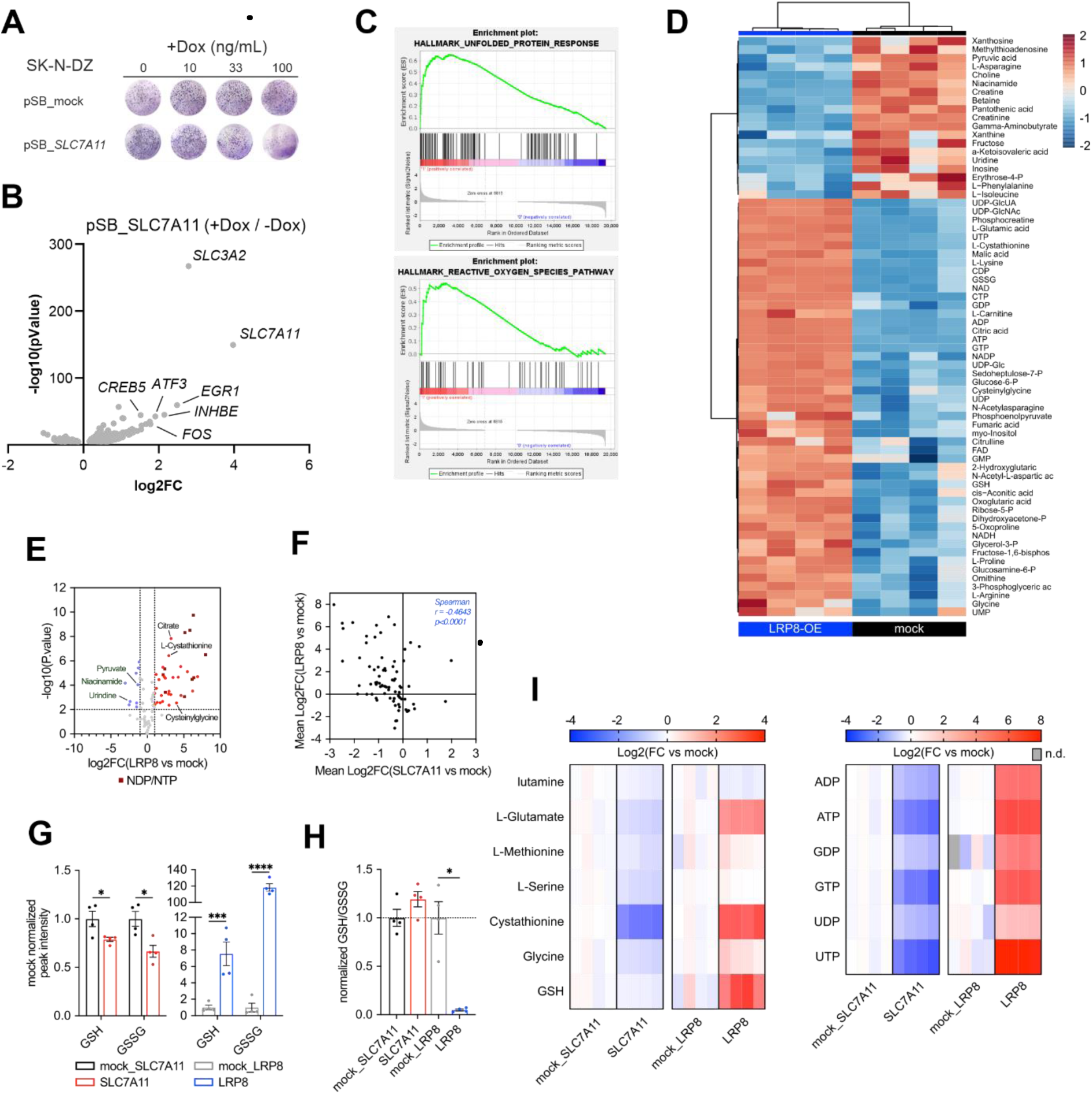
**(A)**. Clonogenic capacity of a clonal SK-N-DZ cell line expressing a Tet-inducible SLC7A11 expression plasmid under different doxycycline concentration. **(B)** Volcano-plot showing differently expressed genes upon SLC7A11 overexpression induced by doxycycline treatment (+Dox) compared to solvent (-Dox). Each dot represents a single gene. Log2 fold-change of mean expression values are plotted against the significance values (log10 transformed adjusted p-values). **(C)** GSEA enrichment plot of the three most differentially enriched gene sets from the HALLMARK collection, namely “unfolded-protein response” and “reactive oxygen species pathway”. **(D)** Heatmap showing 66 metabolites that are significantly different between LRP8-overexpressing SK-N-DZ cells and mock control. Abundance is represented as Log2-transformed normalized peak intensity relative to row mean. **(E)** Volcano plot showing differentially abundant metabolites upon LRP8 overexpression in SK-N-DZ cells compared to mock control. Log2 fold change of mean normalized peak intensities are plotted against the significance values (log10 transformed adjusted p-values). Significantly up-(red) or downregulated (blue) metabolites are indicated and selected metabolites are labelled. NDP/NTP are highlighted in dark red. **(F)** Spearman correlation analysis of Log2 fold changes for 78 polar metabolites compared between LRP8-and SLC7A11-overexpressing SK-N-DZ cells relative to mock controls. **(G)**. Normalized levels of GSH and GSSG in LRP8-and SLC7A11-overexpressing SK-N-DZ cells relative to mock controls. Data are represented as mean ± s.e.m (n=4). **(H)** Relative GSH/GSSG ratio for cells shown in (g). **i**. Heat maps showing log2 fold change of selected metabolites in LRP8-and SLC7A11- overexpressing SK-N-DZ cells relative to mock controls (n=4) (n.d. = not detected). * *P* <0.05, *** *P*<0.005, **** *P*<0.001

**Fig. S7.**
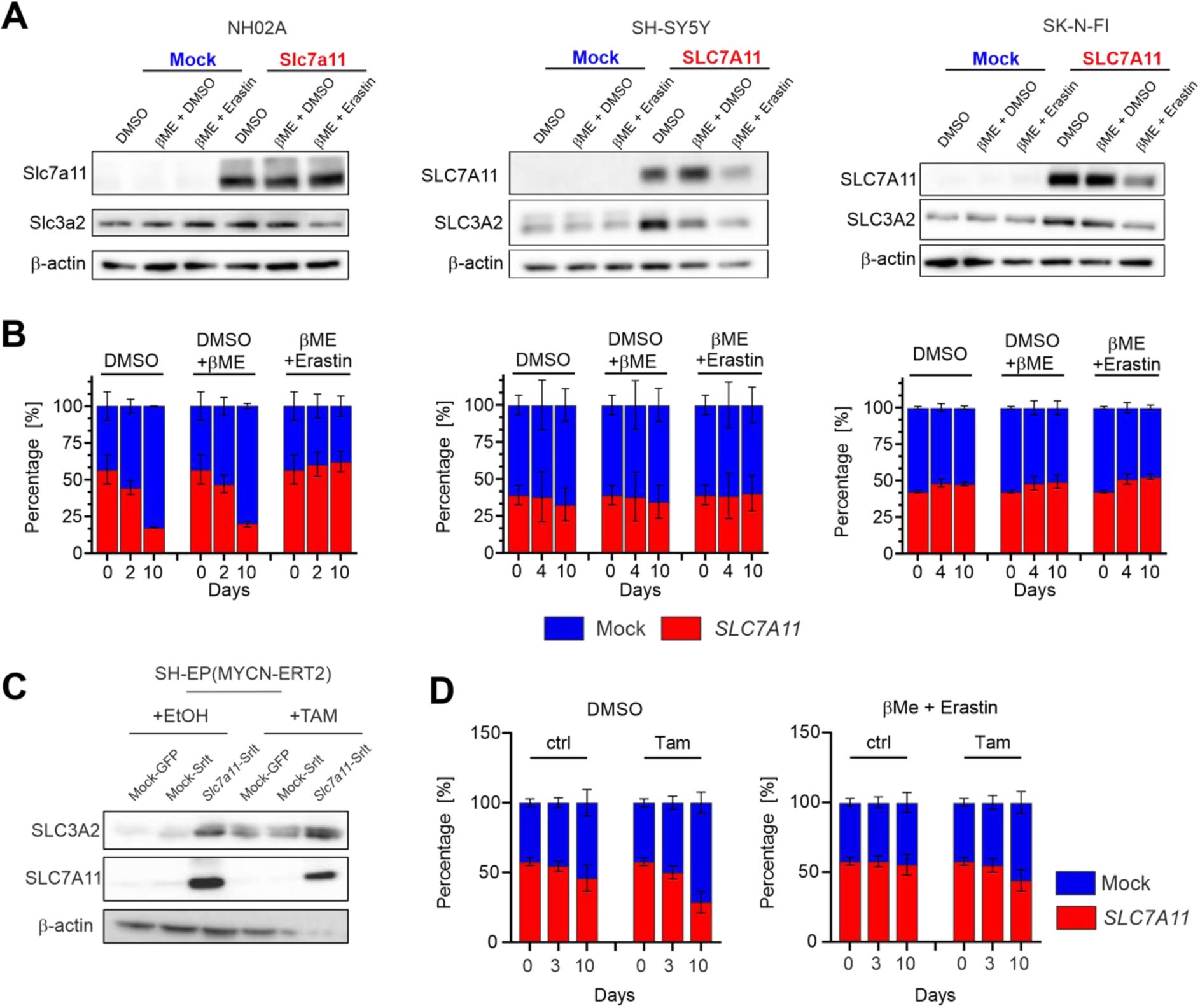
**(A)** Immunoblot analysis of SLC7A11 and SLC3A2 in different neuroblastoma cell lines upon transduction of an SLC7A11 overexpression plasmid (*SLC7A11*^*O*E^) or empty vector (Mock). **(B)** Cell competition assay of a panel of neuroblastoma cell lines overexpressing *SLC7A11* (*SLC7A11*^*O*E^) compared to empty vector (Mock). **(C)** Immunoblot analysis of SLC7A11 and SLC3A2 in a *MYCN-*inducible cell line SH-EP (MYCN-ERT^2^) transduced with an *SLC7A11* overexpression plasmid (*SLC7A11-Srlt*), GFP (Mock-GFP) or empty vector (Mock-Srlt) upon treatment with 0.5 μM tamoxifen or solvent (ethanol) for 24 hours. **(D)** Competition assay of SH-EP (MYCN-ERT^2^) expressing an *SLC7A11* expression plasmid or empty vector (Mock) upon MYCN activation. For the competition assay, cells were seeded at a ratio of 50/50 and 50,000 events were measured by flow cytometry at the depicted time points. Rescue experiments were carried in the presence of βMe alone (50 μM) or a combination of βMe and Erastin (0.5 μM). Time zero in in the different cell lines represents the same experiments ploted repeatidly in order to facilitate visualization. Bars display percentage of eGFP-(blue) and Scarlet-cells (red) with n=3 or means +/-s.e.m. with n=3.

## Methods

### Cell culture

The human neuroblastoma cell line SK-N-DZ was maintained in DMEM medium (Gibco) supplemented with 10% FCS (Gibco), 1X HEPES (Gibco) and 100 U/ml penicillin/streptomycin (Gibco) at 37° C with 5% CO_2_. Cell lines were tested for their identity by SNP genotyping and for mycoplasma contamination by the respective Multiplexion service (Heidelberg, Germany).

### Molecular cloning

The human CRISPR activation pooled library Set A (Addgene plasmid #92379 was a gift from David Root and John Doench) was amplified as described previously (*38*). Briefly, Endura electrocompetent cells (Lucigen, Cat. No. 60242) were used to perform six electroporation reactions on a MicroPulserTM II (BioRad) following the manufacturer’s instructions (pre-set EC1 setting with V: 1.8 kV). Subsequently, cells were pooled and recovered at 37° C for 45 min on a shaking-incubator. The cell suspension was diluted with LB-medium and 2 ml of bacterial suspension was spread evenly on pre-warmed LB-agar plates containing carbenicillin (100 μg/ml, 1x 245 mm square dish per 10,000 gRNAs in library) and incubated for 12 hours at 37° C. The electroporation efficiency was assessed and cells harvested when library representation was >100 bacterial colonies per gRNA. Plasmid DNA was purified using NucleoBond® Xtra Maxi EF (Macherey-Nagel, Cat. No. 740424) according to manufacturer’s instructions. Sequencing libraries were prepared using the NEBNext 2x High-Fidelity Master Mix (NEB, Cat. No. M0541L) with amplification primers that were partially complementary to the lentiviral gRNA backbone with overhangs introducing the Illumina P5/P7 adapter sequences. A pool of staggered P5 primer and 5% PhiX spike-in were used to increase sequence diversity. Maintenance of gRNA distribution and library complexity was confirmed via next-generation sequencing (NGS) on a Hi-Seq2000 with the following read configuration: 125 cycles Read 1, 8 cycles index 1 (i7). Sequencing was performed by the High Throughput Sequencing Unit of the DKFZ Genomics and Proteomics Core Facility.

For the validation and for the scCRISPRa screens, a custom gRNA library was constructed with selected hits from the primary screen. A pool of 82 oligonucleotides containing two gRNAs for each target together with ten NT control gRNAs was ordered from Twist Bioscience (San Francisco, USA) and cloned into a modified CROP-seq-MS2 vector via Golden Gate Assembly (NEB, Cat. No. E5520). The modified CROP-seq-MS2 plasmid was obtained as following: the lentiviral CROP-seq-opti vector (Addgene plasmid #106280 was a gift from Jay Shendure) was sub-cloned via restriction digest of the plasmid with NsiI-HF (NEB, Cat. No. R3127) and SnaBI (NEB, Cat. No. R0130) and the insertion of a synthetic dsDNA fragment coding for a gRNA scaffold sequence with MS2 stem-loop motifs (manufactured by Synbio Technologies, South Brunswick Township, USA). The MS2 loops in the gRNA scaffold allow the recruitment of the p65-HSF1 transactivator complex (expressed from pLentiMPH2). Next, the puromycin resistance gene was extended with a sequence coding for a viral p2A self-cleaving motif and a tagBFP thereby removing the stop-codon from the puromycin resistance gene. For this, the plasmid was linearized with BstEII-HF (NEB, Cat. No. R3162) and MluI-HF (NEB, Cat. No. R3198) and the synthetic dsDNA fragment (manufactured by Twist Bioscience) was inserted to the plasmid backbone thus generating CROP-seq-MS2. The custom gRNA library was amplified in electrocompetent Endura bacteria (Lucigen, Cat. No. 60242) and plasmid DNA purified as described. Library complexity was verified via NGS using the MiSeq V3 kit (Read configuration: 167 cycles Read 1, 8 cycles index 1 (i7)).

Individual gRNAs were cloned into the modified CROP-seq-MS2 or the pXPR_502 (Addgene #96923 was a gift from John Doench and David Root) lentivectors for CRISPRa experiments and the pLKO5_RFP657 backbone for knockout (CIRSPR-KO) experiments via restriction digest of the respective lentivector with BsmBI (NEB, Cat. No. R0739). Oligonucleotides (Sigma-Aldrich) with the gRNA sequences and complementary overhangs were phosphorylated, annealed and inserted into the respective lentiviral delivery vector.

Oligonucleotides used for cloning of individual gRNAs

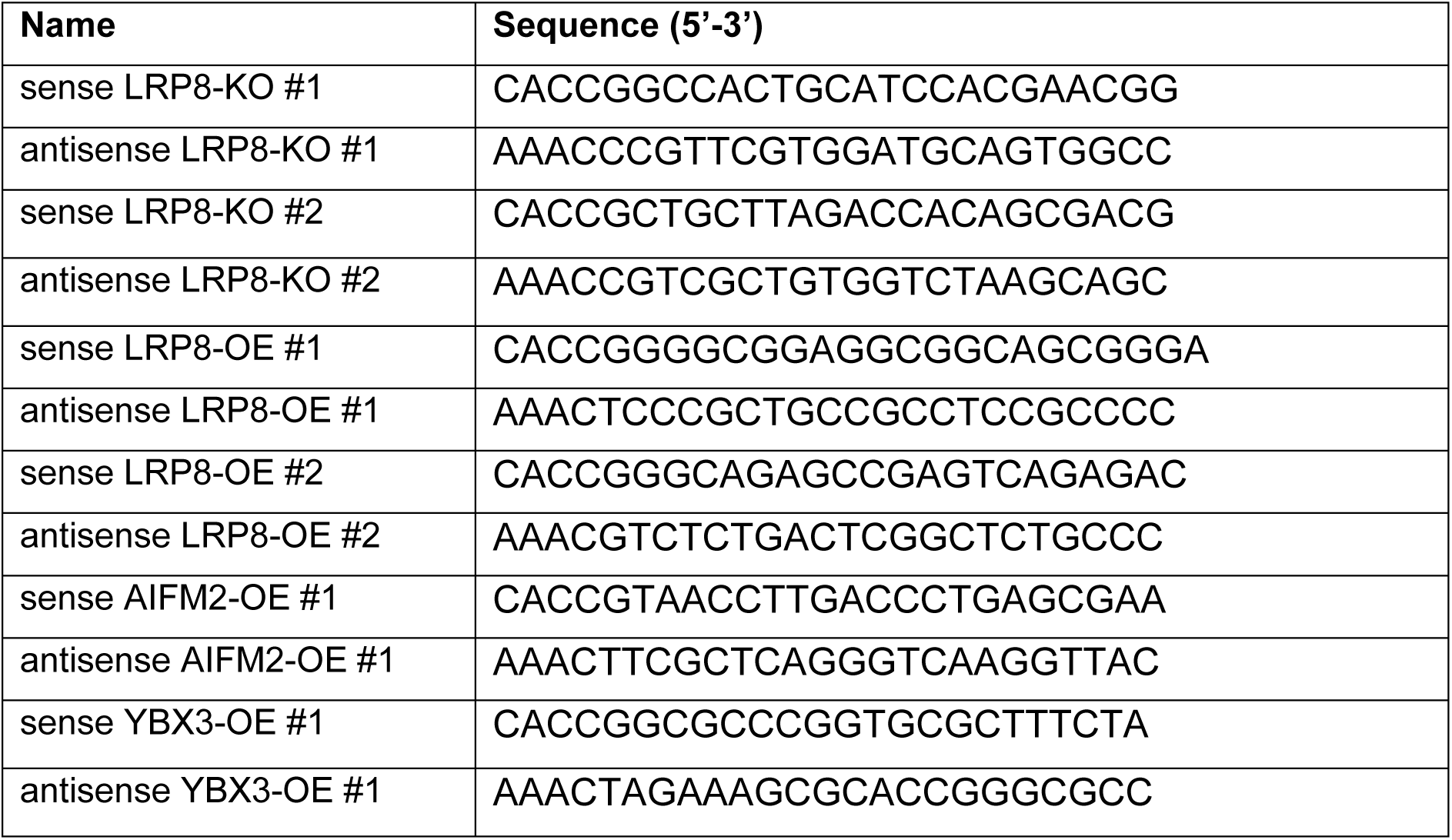
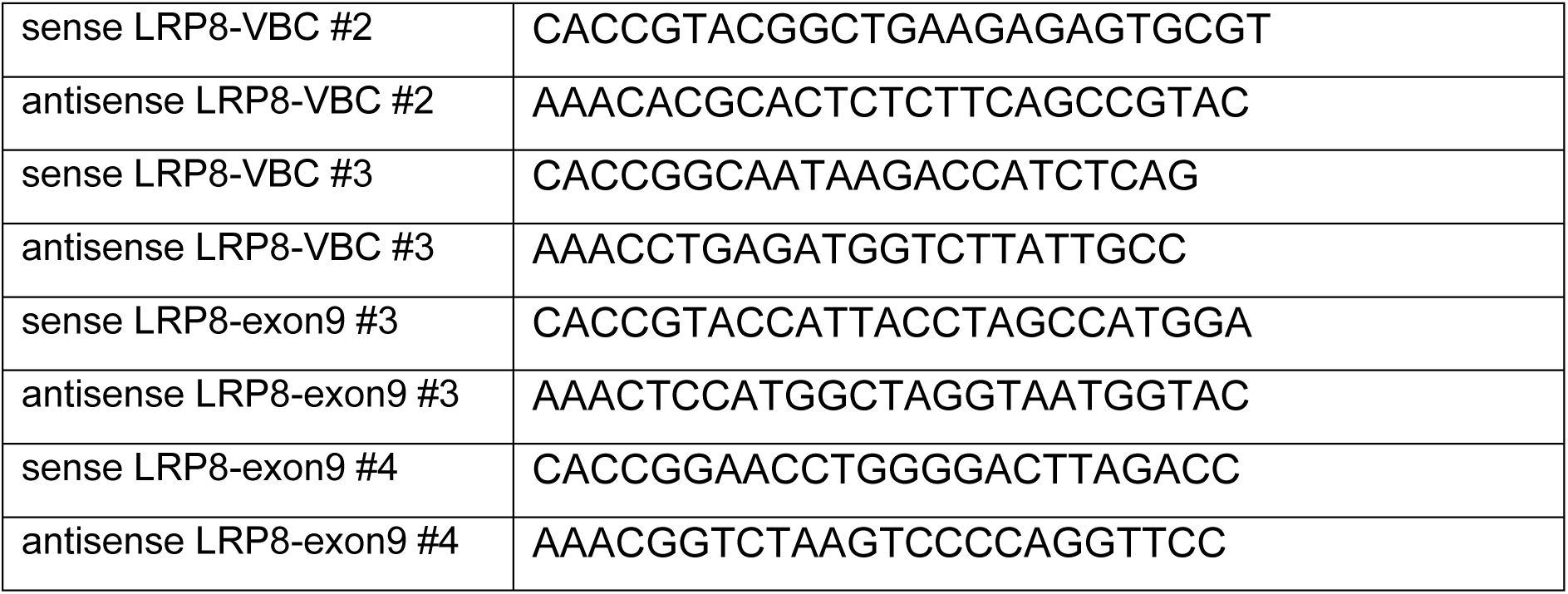

### Lentivirus production

Large-scale lentivirus production was performed using a second-generation lentivirus system and a calcium phosphate transfection kit (Invitrogen, Cat. No. K278001) in HEK293T cells. Briefly, early passaged HEK293T cells were co-transfected with the lentiviral transfer plasmid, a packaging plasmid (psPAX2, Addgene plasmid #12260 was a gift from Didier Trono), as well as with a plasmid encoding the VSV-G envelope (pMD2.G, Addgene plasmid #12259 was a gift from Didier Trono). Viral supernatant was collected 48 hours post transfection, snap frozen and stored at -80° C until use. Alternatively, for the generation of cell expressing the following constructs; p442-LRP8-Flag, p442-Mock, p442-hSLC7A11, p442-mSlc7a11, p442-GFP and p442-mScarlet, HEK293T cells were used to produce replication-incompetent lentiviral particles pseudotyped with the ecotropic envelope protein of the murine leukaemia virus (MLV) or the pantropic envelope protein VSV-G. The third-generation packaging plasmids (MDLg_pRRE and pRSV_Rev) and transfer plasmids were co-transfected into HEK293T cells. Cell cultural supernatant containing viral particle was harvested 48h past transfection and used to transduced cell lines of interest. All experimental procedures for lentivirus production and transduction were performed in a biosafety level 2 laboratory.

### Cell line generation for CRISPRa and CRISPR-KO experiments

Polyclonal SK-N-DZ cells constitutively expressing the CRISPR activation machinery were engineered by transducing wild-type cells with lentiviral particles carrying a dCas9-VP64 (lenti dCas9VP64_Blast, Addgene plasmid #61425 was a gift from Feng Zhang) at a multiplicity of infection (MOI) of ∼0.5. After recovery, cells were selected via Blasticidin treatment (20 μg/ml).

For the validation and scCRISPRa screens, SK-N-DZ cells expressing dCas9-VP64 were transduced with lentiviral particles encoding the p65-HSF1 transactivator complex (pLentiMPH2, Addgene plasmid #89308 was a gift from Feng Zhang). After recovery and selection with hygromycin B (200 μg/ml, Gibco, Cat. No. 10687010), cells were individualized by fluorescence activated cell sorting based on physical parameters (forward scatter and side scatter) and exclusion of dead cells via DAPI staining. Clonal cell lines were established and tested for their CRISPRa potential. Two independent clonal SK-N-DZ cell lines were selected for the validation screen, while the best scoring clonal cell line was used for the scCRISPRa screen.

For the CRISPR-KO experiments, cells were transduced with lentivirus carrying a transgene that encodes a doxycycline-inducible wild-type Cas9 nuclease (pCW-Cas9-EGFP). After recovery, Cas9-inducible cells were transduced with the gRNA lentivirus (pLKO5-RFP657 backbone) at a MOI of ∼0.3. Double positive cells (EGFP/RFP657) were individualized via FACS with the exclusion of dead cells (DAPI negative). Cas9 expression was induced with the addition of doxycycline (1 μg/ml, Sigma-Aldrich, Cat. No. D9891) into the cell culture medium.

### Genome-wide CRISPRa screen to identify negative regulators of ferroptosis

Polyclonal SK-N-DZ cells expressing dCas9-VP64 were transduced with the genome-wide Calabrese CRISPRa library in triplicate at an MOI of ∼0.3. For each replicate, 190 million cells were transduced achieving a representation of ∼1,000 cells per gRNA. After initial recovery for 4 days, cells were selected with puromycin (0.65 μg/ml) for 7 days. For each replicate, cells were split into four groups of which one was harvested to determine baseline gRNA representation, another one maintained as the control group, while two others were treated with one of the ferroptosis inducing agents, RSL3 or Erastin respectively, at a concentration lethal for wild-type SK-N-DZ (300 nM/1 μM). Cells were maintained either with the addition of puromycin (control group) or with the addition of puromycin and the respective drug for additional two weeks. The control cells were passaged maintaining the gRNA representation of 1,000 cells per gRNA throughout the screen. Cells were harvested once the treatment groups reached approximately the same number as the control groups.

The genomic DNA was extracted using a commercial kit (Quick-DNA Midirep Plus kit, Zymo Research, Cat. No. D4075) following the manufacturer’s instructions and NGS libraries prepared as described. Libraries were sequenced as a multiplexed pool on a single lane of a HiSeq2000 chip with the following read configuration (125 cycles Read 1, 8 cycles index 1 (i7)).

### Genome-wide CRISPRa screen analysis and selection of candidate hits

Raw sequencing reads were processed as previously described (*39*). Briefly, reads were trimmed to remove the constant vector sequence upstream of the gRNA sequences with *cutadapt* and then mapped to the reference library with *Bowtie2*. For each gRNA, the assigned reads were counted and normalized (cpm, counts per million reads). Using the control samples, a negative binomial count distribution was estimated and used to determine fold-changes of individual gRNAs in the treatment samples. P-values were computed and corrected for multiple testing (FDR). Finally, a gene-level score was calculated from the mean log fold-change of all gRNAs targeting the same gene. Candidate hits were selected using the top 100 enriched genes and examining protein interactions via StringDB (*40*) network analysis (R package version 2.4.0). Those hits were selected that showed a predicted interaction in the network (total of 36 genes, 162 predicted interactions, p-value: 0.0055).

### Secondary validation CRISPRa screen

Two monoclonal SK-N-DZ cell lines constitutively expressing dCas9-VP64 as well as the p65-HSF1 transactivator complex were transduced with the custom gRNA library targeting the transcription start sites of 36 selected hits from the primary screen (gRNA representation > 2000x). After recovery and selection with puromycin (1 μg/ml), cells were split into one control group and a treatment group with RSL-3 (100 nM). Cells were harvested after two weeks and sequencing libraries prepared as described. Libraries were sequenced as a multiplexed pool on a MiSeq V3 (Read 1: 167 cycles, index 1 (i7): 8 cycles).

### Secondary validation screen analysis

A count matrix of gRNA abundances was determined as described for the primary screen while mapping the reads to the custom reference library. In order to compute enrichment scores and p-values, the robust rank aggregation workflow in the MaGeCK (*41*) pipeline was used.

### Single-cell CRISPRa screen

A monoclonal SK-N-DZ expressing dCas9-VP64 and p65-HSF1 was transduced in duplicate with the custom gRNA library in the CROP-seq-MS2 backbone at a low MOI of ∼0.07 to minimize the integration of multiple gRNAs. Transduction efficiency was confirmed via flow cytometric analysis of tagBFP expression. Cells were selected with puromycin at 1 μg/ml 24 hours post-transduction. Puromycin selection was continued for additional 4 days at 1.5 μg/ml. Ten days after transduction, cells were collected and ∼20,000 individual cells per lane were partitioned by the 10X Chromium Controller. Single-cell RNA-seq libraries were constructed using the Chromium Single Cell Gene Expression v3.1 kit (10X Genomics, Cat. No. 1000128) following the manufacturer’s protocol thereby retaining 40 ng of full-length cDNA for the enrichment of gRNAs. Guide RNA sequences were selectively amplified adapting a hemi-nested PCR approach that was previously described (*42*) using the gRNA_enrichment1_fw (5’-GTGACTGGAGTTCAGACGTGTGCTCTTCCGATCTCTTGTG-GAAAGGACGAAACACCG-3’) and gRNA_enrichment1_rv (5’-CTACACGAC-GCTCTTCCGATCT-3’) primers and the 2x KAPA HiFi HotStart Ready Mix (Roche, Cat. No. KK2601). In a subsequent PCR, 1 ng of purified PCR product was used as input to construct sequencing ready libraries with the gRNA-enrichment2_fw (5’- CAAGCAGAAGACGGCATACGAGAT[index1]GTGACTGGAGTTCAG-3’) and gRNA_enrichment2_rv (5’-AATGATACGGCGAC-CACCGAGATCTACACTCTTTCCCTACACGACGCTC-3’) primers. The whole-transcriptome single-cell libraries were sequenced on a NovaSeq6000 with 28 cycles Read 1, 8 cycles index 1 (i7) and 91 cycles Read 2. The gRNA enrichment libraries were sequenced on a NextSeq550 high with the following read configuration: 32 cycles Read 1, 8 cycles index 1 (i7) and 120 cycles Read 2.

### Single-cell CRISPRa screen analysis

Whole-transcriptome single-cell reads were mapped to the reference genome (GRCh 38) and count matrices were generated using Cell Ranger (10X Genomics, version 6.0.1) with default parameters. Single-cell sequencing data from two individual 10X Chromium lanes were combined using the aggregation function (cellranger aggr). Data normalization and downstream analysis were performed with the R package Seurat (version 4.0.1). Based on the inflection point per cell barcode, a threshold of 1,000 detected features per cell was set to filter out empty droplets as well as cells with low complexity. The count matrix was filtered for genes detected in at least 10 cells. For each cell, UMI counts of each gene were divided by the total number of counts and multiplied by a scaling factor (10,000) after which they were log-transformed. Guide RNA enrichment libraries were processed via fba (*43*) (version 0.0.11) and mapped to the custom gRNA library thereby allowing two mismatches. The Seurat object was filtered for cells that were assigned to a particular gRNA and cells with a strong CRISPRa phenotype were determined using Seurat’s Mixscape function. Differential gene expression analysis was performed for each CRISPRa group with DESeq2 (version 1.32.0) using cells assigned to the NT group as control. The top 50 differentially expressed genes were selected (ranked by the adjusted p-value) from each overexpression group and merged together to create the gene expression signature list. For Fig. 1F, functionally related differentially expressed genes were labelled based on overrepresented terms that were retrieved from the Gene Ontology, the Reactome, the KEGG and the Molecular Signatures databases and based on enrichment analysis performed with the R package clusterProfiler (version 4.0.2).

### SLC focused CRISPR based screens for selenium relevant transporters

SK-N-DZ were transduced with the SLC-focused CRISPR based library in duplicate at an MOI of 0.5. For each replicate, 10 million cells were transduced to achieve a representation of >1000. After recovery for 3 days, the cells were selected with 0.5 μg/ml puromycin and were maintained with 500nM Lip-1 for 5 days. After selection, for each replicate, the cells were split into five groups which were treated with 20nM Na2SeO3, 20nM Selenocystiene, 200nM Selenomethione, 50μM ßME and 500nM Lip-1 respectively. The cells were passaged maintaining representation >1000 cells per gRNA throughout the screen. Cell pellets were harvested at day 14 of treatment. The genomic DNA was extracted using a commercial kit (QIAGEN-DNeasy) following manufacturer’s instructions. Libraries were sequenced as a multiplexed pool on a Nextseq500 (8 cycles index 2 (i5), 8 cycles index 1 (i7), 75 cycles Read 1, 75 cycles Read 2)

### Generation of *LRP8*-knockout cell lines

SK-N-DZ cells were transduced with lentiviral particles encoding LRP8 gRNAs (lentiCRISPRv2 blast, Addgene plasmid #98293 was a gift from Brett Stringer). After recovery and selection with blasticidin (10μg/ml), cells were subcloned via limiting dilution. Clonal cell lines were established and tested for LRP8 expression by Western Blot. Two clonal cell lines were selected for further experiments.

Clonal HT1080 and A375 cells constitutively expressing Cas9 were transfected with plasmids encoding gRNAs (pKLV2-U6gRNA5(BbsI)-PGKpuro2AmAG-W, Addgene plasmid #67976 was a gift from Kosuke Yusa) targeting introns flanking *LRP8* exon 9. After selection with puromycin (1μg/ml), cells were grown by limiting dilution. Clonal cell lines were established and tested by genotyping PCR and Western Blot. Two clonal cell lines were selected for further experiments and subsequent CRISPR based screens.

### SLC focused CRISPR based screens in *LRP*-KO and wild-type

*LRP8*-KO and WT cell lines were transduced with the SLC focused CRISPR based library at an MOI of 0.5. 10 million cells for each cell lines were transduced achieving >1000 cells per gRNA. After recovery and selection with 1μg/ml puromycin, the *LRP8*-KO and -WT cell lines were split and maintained at the gRNA representation of >1000. Cell pellets were harvested after 2 weeks of passaging. The genomic DNA was extracted with a commercial kit (QIAGEN-DNeasy) following the manufacturer’s instructions. Libraries were multiplexed and sequenced on a Nextseq 500 (8 cycles index 2 (i5), 8 cycles index 1 (i7), 75 cycles Read 1, 75 cycles Read 2). The mapping of raw sequencing reads to the reference library and computing of enrichment scores and P-values were processed using the MaGeCK pipeline. The MaGeCKFlute was used for identification of gene hits and associate pathways.

### Cell lines generation for competition assay

Human neuroblastoma cell lines (SK-N-DZ, SK-N-FI, SH-SY5Y and SH-EP MYCN-ER) and the murine neuroblastoma cell line (NH02A) expressing a lentiviral construct carrying *SLC7A11* cDNA or an empty vector were generated using the methods described above. Briefly, lentiviral constructs were transduced into the corresponding cell line labeled with mScarlet or eGFP. The combination of construct/fluorophore is described in the figures present in the main text. Established cell lines were counted and mixed at a 1:1 ratio and the distribution was monitored during the period indicated in the main text. Rescue experiments were carried using a combination of ßMe (50 μM) and Erastin (0.5 μM).

### Analysis of Cell Viability

The impact of various compounds on cell viability was analyzed using the CellTiter-Glo (CTG) assay (Promega). To determine changes in cell viability, 3000 cells were seeded in full medium in 96-well plates (Greiner Bio One) 24 h prior to the treatment. Cells were then treated for 72 h with the indicated concentration of compounds. Cell viability was analyzed using the CTG assay following the manufacturer’s instructions. For in vitro clonogenic assays, 200 cells were seeded in 12-well plate for 14 days with each particular experimental condition, and colonies were stained with 1 mL 0.01% (w/v) crystal violet.

### Analysis of Lipid peroxidation

Approximately 10^5^ cells were seeded in six-well plates. Before lipid peroxidation being analyzed, medium was removed, C11-BODIPY (wave length: 581/591 nm) diluted in Hank’s Balanced Salt Solution (HBSS; Gibco) and added to wells at a final concentration of 4 μM. After 15 min staining at 37° C inside the tissue culture incubator, cells were harvested gently and levels of lipid peroxidation immediately analyzed using a BD FACS Aria™ III cell sorter.

### Transient siRNA-mediated gene knockdown

SK-N-DZ cells were seeded in 12-well plates (200,000 cells/well) and 24h later transiently transfected with a mix of RNAiMax (0.04 μl/well; Thermo Fisher Scientific) and 0.01 μM/well of siRNA following the manufacturers’ instructions.

### Metabolic Profiling

Water-soluble metabolites from *SLC7A11*-overexpressing cells were extracted with 0.5 ml ice-cold MeOH/H2O (80/20, v/v) containing 0.01 μM lamivudine (Sigma-Aldrich) and water-soluble metabolites from *LRP8*-overexpressing cells were extracted with 1.0 ml ice-cold MeOH/CH3CN/H2O (50/30/20, v/v/v) containing 4 μM of the internal standards D4-glutaric acid and D8-phenylalanine. The suspensions were vigorously mixed, sonicated, mixed again and centrifuged for 5 min at 13,000g. After centrifugation of the resulting homogenates, supernatants were transferred to a RP18 SPE column (Phenomenex) that had been activated with 1.0 ml CH3CN and conditioned with 1.0 mL of MeOH/H2O (80/20, v/v). The remaining cell pellet was again resuspended in 0.5 mL ice-cold MeOH/CH3CN/H2O (50/30/20, v/v/v), mixed, sonicated and centrifuged as before. The supernatant was transferred to the same column again and the eluate was collected in the same tube as before. The eluate was dried in a centrifugal evaporator Savant (Thermo Scientific) or SpeedVac (Labconco) and dissolved in 5 mM NH4OAc in CH3CN / H2O (50/50, v/v) in 50μL for *SLC7A11*-overexpressing cells and in 100μL/1*10^6 cells for *LRP8*-overexpressing cells.

For LC-MS analysis of water-soluble metabolites from *SLC7A11*-overexpressing cells, 3 μl of each sample was applied to a ZIC-cHILIC column (Sigma-Aldrich SeQuant ZIC-cHILIC, 3μm particle size, 100*2.1 mm). Metabolites were separated at 30°C by LC using a DIONEX Ultimate 3000 UPLC system (Thermo Fisher Scientific) and the following solvents: Solvent A consisting of 5 mM NH4OAc in CH3CN/H2O (5/95, v/v) and solvent B consisting of 5 mM NH4OAc in CH3CN/H2O (95/5, v/v). At a flow rate of 200 μl/min, a linear gradient starting at 100% solvent B decreasing to 40% solvent B over 23 min was applied followed by 17 min constant elution with 40% solvent B, followed by a linear increase to 100% solvent B over 1 min. Recalibration of the column was achieved by 7 min pre-run with 100% solvent B. MS-analyses were performed on a high-resolution Q-Exactive mass spectrometer (Thermo Fisher Scientific) in alternating positive-and negative full MS mode applying the following scan and HESI source parameters: Scan Range: 69.0 - 1000 m/z. Resolution: 70,000, AGC-Target: 3E6, Maximum Injection Time: 200 ms. Sheath gas: 30, auxiliary gas: 10, sweep gas: 3, Aux Gas Heater temperature: 120 °C. Spray voltage: 2.5 kV (pos)/3.6 kV (neg), capillary temperature: 320 °C, S-lens RF level: 55.0.

For LC/MS analysis of water-soluble metabolites from *LRP8*-overexpressing cells, 3 μL of sample was applied to an Amid-HILIC column (Thermo Fisher Scientific Accucore 150 Amid-HILIC, 2.6μm particle size, 100 × 2.1 mm). Metabolites were separated at 30°C by LC using a DIONEX Ultimate 3000 UPLC system (Thermo Fisher Scientific) and the following solvents: Solvent A consisting of 5 mM NH4OAc in CH3CN/H2O (5/95, v/v) and solvent B consisting of 5 mM NH4OAc in CH3CN/H2O (95/5, v/v). At a flow rate of 350 μL/min, a linear gradient starting at 98% solvent B for 1 min, followed by a linear decrease to 40% solvent B within 5 min, then maintaining 40% solvent B for 13 minutes, then returning to 98% solvent B within 1 minute. The column was equilibrated at 98% solvent B for 5 minutes prior every injection. The eluent was directed to the ESI source of the MS instrument from 0.5 minutes to 19 minutes after sample injection. MS analysis was performed on a high-resolution Q-Exactive plus mass spectrometer (Thermo Fisher Scientific) in alternating positive-and negative full MS mode applying the following scan and HESI source parameters: 69-1000 m/z; resolution: 70,000; AGC-Target: 3E6; maximum injection time: 50 ms; sheath gas: 30; auxiliary gas: 10; aux gas heater temperature: 120°C; spray voltage: 3.6 kV (pos)/2.5 kV (neg), capillary temperature: 320°C, S-lens RF level: 55.0. For the fragmentation of water-soluble metabolites, the following ddMS2 settings were applied in both modes: Resolution: 17,500; AGC-Target: 1E5; maximum injection time: 50 ms; Loop count: 1; CE: 20, 50, 80; Apex trigger: 0.1 to 10s; minimum AGC target: 2E3; Dynamic exclusion: 20 s.

Signal determination and quantitation was performed using TraceFinder™ Software Version 3.3 (Thermo Fisher) or using El-Maven Software Version 0.12.0 (https://elucidata.io/el-maven/).

### Analysis of oxygen consumption rate

Oxygen consumption rate (OCR) of SK-N-DZ (*SLC7A11*-overexpressing and Mock) cells was measured using the Seahorse XF96 Analyzer (Seahorse Biosciences) and the Seahorse XF Cell Mito Stress Test Kit with oligomycin (final concentration 1.5 μM), followed by FCCP (final concentration 2 μM), and Rotenone (final concentration 0.5 μM). For this experiment 40,000 cells/well were plated a day before the experiment. This allowed for a 60–70% confluency at the time of measurement. Data were normalized to total protein amount (Bio-Rad protein assay).

### Selenium speciation analyses

We measured total selenium by inductively coupled plasma sectorfield mass spectrometry (ICP-sf-MS) and the selenium species selenite (Se-IV), selenate (Se-VI), selenomethionine-bound selenium (Se-MET), selenocystine-bound selenium (Se-Cys), thioredoxin reductase-bound selenium (Se-TrxR), glutathione-peroxidase-bound selenium (Se-GPx), selenoprotein-P-bound selenium (SELENOP) and albumin-bound selenium (Se-HSA) using ion exchange chromatography (IEC) coupled with ICP-dynamic reaction cell mass spectrometry (ICP-DRC-MS) in analogy to methodologies previously established (*44*). The experimental settings for ICP-sf-MS (ELEMENT II, Thermo Scientific, Bremen Germany) were: radio frequency power: 1260 W, plasma gas flow: 16L Ar/min auxiliary gas flow: 0.85L Ar/min, nebulizer gas flow: 1.085 L Ar/min, daily optimized, dwell time 300 ms, ions montored: 77Se, 78Se, high resolution mode. For speciation of selenium compounds, we used the hyphenated system from Perkin Elmer (Rodgau, Germany) comprising of a NexSAR gradient HPLC pump, autosampler and NexION 300 D ICP-DRC-MS, completely controlled by Clarity software. The separation column for species separation was an ion exchange pre- and analytical column AG-11+AS- 11 (250 × 4 mm I.D.) from Thermo Dionex (Idstein, Germany). The sample volume was 50 μl. The mobile phases and chromatographic gradient were previously published(*44*). Briefly, the flow rate was 0.80 ml/min. The experimental settings for ICP-DRC-MS) were: radio frequency power: 1250 W, plasma gas flow: 15L Ar/min auxiliary gas flow: 1.05L Ar/min, nebulizer gas flow: 0.92 L Ar/min, daily optimized, dwell time 300 ms, ions montired: 77Se, 78Se, 80Se, DRC reaction gas: CH4 reaction at 0.58 ml/min, DRC rejection parameter q: 0.6. Five-point calibration curves from 0-5000 ng/L were linear with r^2^ for the three Se isotopes being better than 0.999881. Data files from selenium chromatograms were processed with Clarity software for peak area integration.

### Analyses of pediatric cohorts of neuroblastoma samples

RNA sequencing of 498 neuroblastoma cases was performed as described previously (*45*). In short, mRNA purification was done using the Dynabeads mRNA Purification Kit (Invitrogen) and library construction was performed according to the standard TruSeq protocol. Clusters were generated according to the TruSeq PE cluster Kit version 3 reagent preparation guide (for cBot-HiSeq/HiScanSQ). Paired-end sequencing with 100bp read length was performed on the Illumina HiSeq 2000 platform. Raw data processing, read mapping, and gene expression quantification were done using the Magic-AceView analysis pipeline and AceView transcriptome reference (http://www.aceview.org) as described previously(*45*). Genes with generally low read counts were removed using R (v4.1.1) and the function ‘filterByExpr’ in R-package edgeR (v3.34.1). Differential gene expression analysis was done using the empirical Bayes method implemented in the R-package limma (v3.48.3).

### *In vivo* orthotopic mouse experiments

All studies involving mice and experimental protocols were conducted in compliance with the German Cancer Center Institute guidelines and approved by the governmental review board of the state of Baden-Wuerttemberg, Regierungspraesidium Karlsruhe, under the authorization number G-176/19, followed the German legal regulations. Mouse strains used in the study: NOD.Cg-Prkdc^scid^Il2rgtm1^Wjl^/SzJ (NSG, JAX stock #005557). Female mice, 3 – 4 months of age, were used for experiments. Mice were housed in individually ventilated cages under temperature and humidity control. Cages contained an enriched environment with bedding material. To generate orthotopic mouse models for neuroblastoma, 2×10^5^ SK-N-DZ cells were transplanted into the right adrenal gland after surgical site was prepared. Cells were resuspended in a 1:1 (vol/vol) mix of growth factor-reduced matrigel (Corning) and PBS. Overall, 20 μl of this cell suspension was injected into the right adrenal gland of anaesthetized mice. After tumor cell transplantation, we monitored the mice for evidence of tumor development by bioluminescent signal using an IVIS Spectrum Xenogen device (Caliper Life Sciences). We observed a clear signal from the tumors one week after the injection of 2×10^5^ SK-N-DZ cells. For liproxstatin (Lip-1) we used 10 mg/kg/d for the first 5 days. Lip-1 treatment (every second day) continued in one group for another 2 weeks. Animals’ health was monitored daily and mice were euthanized as soon as they reached abortion criteria defined in the procedure. Sample size was calculated with the help of a biostatistician using R version 3.4.0. Assumptions for power analysis were as follows: *α* error, 5%; *β* error, 20%. Mice were randomized into treatment groups prior to treatment. In case animals had to be sacrificed before the pre-defined endpoint (due to weight loss or other termination criteria), they were excluded from any downstream analyses. All animal experiments were blinded during experiments and follow up assessment.

## Notes

**Competing interests:** The authors declare the following financial interests/personal relationships which may be considered as potential competing interests: M.C. is the co-founder of ROSCUE THERAPEUTICS GmbH and author of patent application related to ferroptosis. G.S-F. is co-author of patent applications related to SLCs, co-founder of Solgate GmbH as well as the Academic Project Coordinator of the IMI grants RESOLUTE and Resolution in partnership with Pfizer, Novartis, Bayer, Sanofi, Boehringer-Ingelheim and Vifor Pharma. The G.S-F. laboratory receives funds from Pfizer. All other authors declare no other relevant conflicts of interest.

### Competing Interest Statement

Competing interests: The authors declare the following financial interests/personal relationships which may be considered as potential competing interests: M.C. is the co-founder of ROSCUE THERAPEUTICS GmbH and author of patent application related to ferroptosis. G.S-F. is co-author of patent applications related to SLCs, co-founder of Solgate GmbH as well as the Academic Project Coordinator of the IMI grants RESOLUTE and Resolution in partnership with Pfizer, Novartis, Bayer, Sanofi, Boehringer-Ingelheim and Vifor Pharma. The G.S-F. laboratory receives funds from Pfizer. All other authors declare no other relevant conflicts of interest.

